# Mechanosensing through direct binding of tensed F-actin by LIM domains

**DOI:** 10.1101/2020.03.06.979245

**Authors:** Xiaoyu Sun, Donovan Y. Z. Phua, Lucas Axiotakis, Mark A. Smith, Elizabeth Blankman, Rui Gong, Robert C. Cail, Santiago Espinosa de Los Reyes, Mary C. Beckerle, Clare M. Waterman, Gregory M. Alushin

## Abstract

Mechanical signals transmitted through the cytoplasmic actin cytoskeleton must be relayed to the nucleus to control gene expression. LIM domains are protein-protein interaction modules found in cytoskeletal proteins and transcriptional regulators; however, it is unclear if there is a direct link between these two functions. Here we identify three LIM protein families (zyxin, paxillin, and FHL) whose members preferentially localize to the actin cytoskeleton in mechanically-stimulated cells through their tandem LIM domains. A minimal actin-myosin reconstitution system reveals that representatives of all three families directly bind F-actin only in the presence of mechanical force. Point mutations at a site conserved in each LIM domain of these proteins selectively disrupt tensed F-actin binding *in vitro* and cytoskeletal localization in cells, demonstrating a common, avidity-based mechanism. Finally, we find that binding to tensed F-actin in the cytoplasm excludes the cancer-associated transcriptional co-activator FHL2 from the nucleus in stiff microenvironments. This establishes direct force-activated F-actin binding by FHL2 as a mechanosensing mechanism. Our studies suggest that force-dependent sequestration of LIM proteins on the actin cytoskeleton could be a general mechanism for controlling nuclear localization to effect mechanical signaling.

## Introduction

The sensing of mechanical stimuli by cells plays a pivotal role in maintaining tissue homeostasis^1^, and defects in mechanical signal transduction (“mechanotransduction”) are associated with numerous diseases, including tumorigenesis^2^, cancer metastasis^3^, and developmental disorders^4^. Cells probe the mechanical properties of their microenvironments through integrin-mediated focal adhesions (FAs) linked to actomyosin stress fibers (SFs)^5, 6^. Forces transmitted through these cytoskeletal networks are transduced into gene expression programs by the downstream nuclear localization of transcriptional co-activators^7, 8^. The upstream mechanically-regulated binding interactions between these factors and cytoskeletal partners which govern their nuclear localization are unknown.

The transcriptional co-activator FHL2 (Four-and-a-Half LIM domains 2, also known as DRAL and SLIM-3) is dysregulated in a wide variety of cancers^9–11^, cardiomyopathies^12, 13^, and developmental disorders^14, 15^. FHL2 has been reported to mediate mechanotransduction via enhanced nuclear localization in soft environments^16^. It is composed solely of LIM domains, protein-protein interaction modules^17^ that confer specific protein-binding activities to diverse actin cytoskeleton-associated proteins^18^ and homeobox transcription factors^19^. Many LIM-domain containing proteins exhibit diminished FA / SF recruitment when actomyosin contractility is pharmacologically suppressed^20, 21^, suggesting that LIM domains could serve as a mechanical response module. LIM-domain dependent cytoskeletal “mechanoaccumulation” has been well-established for the SF repair factor zyxin, defined as recruitment to SFs in response to uniaxial cyclic stretch^22^, as well as rapid localization to micron-scale regions along SF known as “strain sites” formed spontaneously in contractile cells^23^. Selected additional members of the paxillin families of FA / SF proteins have also been reported to mechanoaccumulate^24, 25^, but it is unknown how widely distributed this activity is in the LIM protein superfamily. Although direct actin-binding has been speculated to play a role in mechanoaccumulation^18, 26^, zyxin exhibits negligible association with F-actin in co-sedimentation assays^27^, and a force-dependent molecular mark in SFs recognized by LIM domains remains unknown.

Here we show that mechanoresponder LIM domains directly bind tensed F-actin through a conserved mechanism. We find that tensed F-actin binding serves as the dominant mechanism to retain FHL2 in the cytoplasm in stiff environments. The degree of FHL2 nuclear localization is quantitatively related to the availability of cytoskeletal binding sites and its capacity to engage them, supporting a simple model where tensed F-actin acts as a cytoplasmic sink for retaining FHL2 in stiff environments to prevent its nuclear shuttling. Our finding that force-activated F-actin binding is widely distributed in the LIM protein superfamily suggests that it could be a general mechanism for coordinating mechanotransduction through the cytoskeleton.

## Results

### LIM domains from three protein families are sufficient for mechanoaccumulation, which can be negatively regulated by sequence context

To identify mechanoresponsive LIM proteins, we screened a curated list of 28 eGFP-tagged LIM proteins (Fig. 1a) identified in FA / SF proteomics studies^20, 21^ for enhanced localization to the actin cytoskeleton of mouse embryonic fibroblasts (MEFs) mechanically stimulated by uniaxial cyclic stretch (Methods, Extended Data Fig. 1a, Extended Data Fig. 2, 3). We found members of the FHL (FHL2 and FHL3), paxillin (HIC5), zyxin (zyxin, FBLIM1, and TRIP6), and ALP (PDLIM1 and PDLIM2) families exhibited significant increases in actin enrichment in stretched cells versus unstretched controls (Extended Data Fig. 2, 3). To investigate which families of mechanoresponsive LIM proteins identified in our screen could feasibly mechanoaccumulate through a common molecular mechanism, we used live-cell confocal imaging to monitor the association rate of representative mechanoresponsive proteins to spontaneously generated SF strain sites marked by zyxin-fusionRed in a human osteosarcoma cell line (U2OS, Extended Data Fig. 4) plated on glass (Methods). We found that paxillin family member HIC5, zyxin family member FBLIM1, and FHL family members FHL2 and FHL3 are recruited to strain sites with kinetics that are indistinguishable from zyxin within our 10 s frame rate of imaging, as has previously been reported for paxillin (Extended Data Fig. 4c)^24, 25^. Their coherent binding kinetics suggest that these LIM proteins likely recognize a common molecular mark at strain sites. In contrast, an ALP family mechanoresponder, PDLIM1, accumulates at strain sites at a substantially later time point comparable to α-actinin (Extended Data Fig. 4c), with which ALP/Enigma family LIM proteins are known to directly interact^28, 29^. As this suggests ALP family members mechanoaccumulate by an idiosyncratic molecular mechanism, we focused on the FHL, paxillin, and zyxin families for the remainder of our study.

**Fig. 1.**
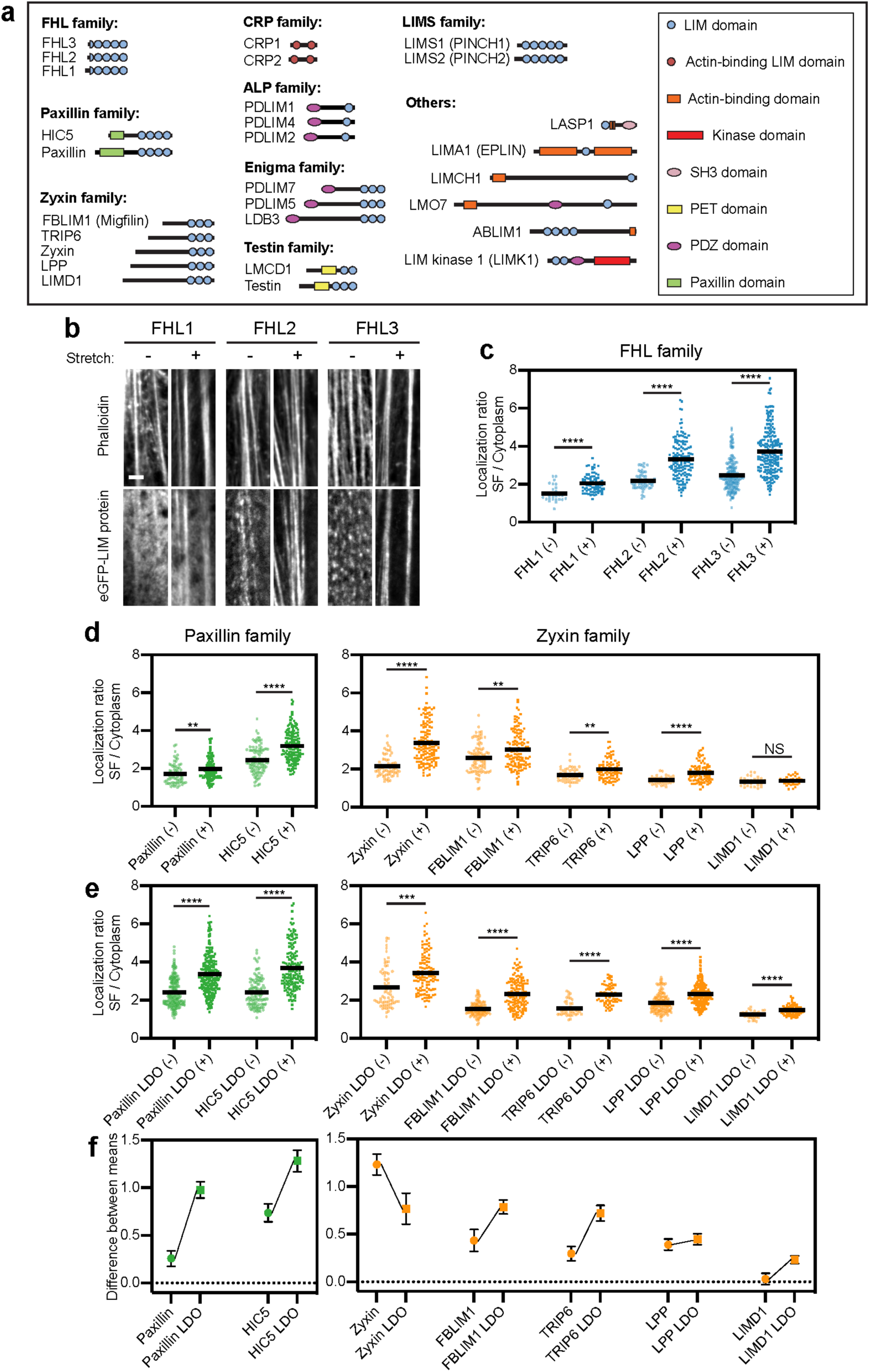
**The FHL, paxillin, and zyxin families of LIM-domain proteins accumulate on SFs in response to mechanical stimulation. a**, Cartoon schematics of LIM-domain proteins examined in this study, drawn to scale by primary sequence. **b**, Confocal slices of phalloidin-stained SFs (top) and eGFP-labeled FHL-family proteins (bottom). Images are cropped around individual SFs. Scale bar, 2 μm. **c**, SF enrichments of eGFP-labeled FHL-family proteins in unstretched (-) and stretched (+) MEFs. Each data point is obtained from a single SF (28 ≤ n ≤ 226, N = 2 biological replicates). **d**,**e**, SF enrichments of FL (**d**) and LDO (**e**) proteins of paxillin and zyxin families in unstretched (-) and stretched (+) MEFs. Each data point is obtained from a single SF (27 ≤ n ≤ 243, N = 2 biological replicates). Bars represent means. Games-Howell’s multiple comparison test after Welch’s ANOVA. NS, p > 0.05; ** p < 0.01; *** p < 0.001; **** p < 0.0001. Outliers (Methods, Extended Data Table 1) are not displayed. **f**, Difference between mean SF enrichments in stretched and unstretched MEFs of FL (circles) versus LDO (squares) versions of each protein. Error bars represent standard error of the difference between the means.

Hypothesizing that non-SF actin networks, which are not anchored to the substrate through FAs^30^, could mask weak mechanoaccumulation of some LIM proteins, we re-assayed all zyxin, paxillin, and FHL family members for stretch-induced recruitment specifically to substrate-anchored SFs in MEFs (Methods, Extended Data Fig. 1b). With the exception of the zyxin-family protein LIMD1, all the LIM proteins from these three families significantly mechanoaccumulate on SFs (Fig. 1b-d, Extended Data Fig. 5a), including endogenous FHL2 (Extended Data Fig. 5b). We next investigated whether the LIM domains from these proteins are sufficient for mechanoaccumulation, as has previously been reported for paxillin^24, 25^ and zyxin^22, 23^. We generated eGFP-tagged LIM-domain-only (LDO) versions of the paxillin- and zyxin-family proteins (FHL proteins only contain LIM domains, Fig. 1a), which revealed that all of the LDOs, including LIMD1 LDO, mechanoaccumulate on SFs (Extended Data Fig. 5a, Fig. 1e). Furthermore, with the exception of the zyxin LDO, all LDOs exhibit enhanced SF enrichment compared to their full-length counterparts (Fig. 1f). This suggests that while non-LIM sequence has the capacity to negatively regulate LIM domains *in cis* through autoinhibition and/or sequestration at other cellular compartments such as FAs^19^, mechanoaccumulation is an intrinsic property of mechanoresponder LIM domains.

### LIM domains are functionally swappable mechanoresponder modules

To test whether cytoskeletal mechanoaccumulation by LIM domains is a modular activity, we examined the capacity of FHL2/3 to functionally substitute for the zyxin LIM domains. Zyxin’s primary sequence has previously been established to have functionally separable regions, with its N-terminal region featuring motifs for binding α-actinin and VASP to facilitate actin repair, and its 3 C-terminal LIM domains mediating mechanoaccumulation^22, 23^ (Fig. 2a). We generated zyxin-FHL2 and zyxin-FHL3 chimeras by replacing the zyxin LIM domains with the coding sequences of FHL2/3 (Fig. 2a), then assayed them for functional rescue in zyxin null MEFs previously reported to feature actin repair defects^23, 31^ (Fig. 2).

**Fig. 2.**
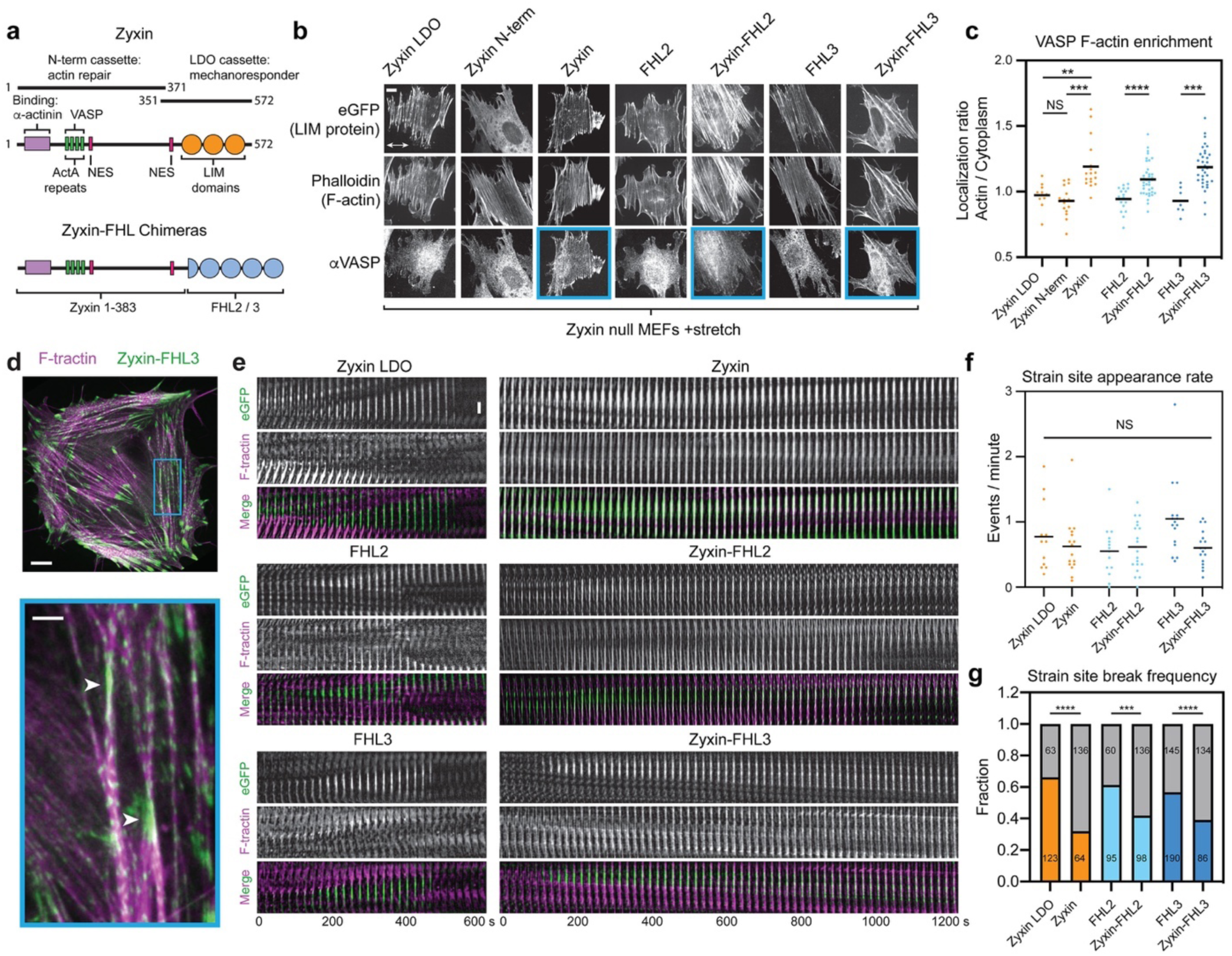
**The modular mechanoresponder LIM domains of FHL confer actin repair activity when fused to the zyxin N-terminus. a,** Schematic of zyxin and zyxin-FHL chimeras; residue ranges comprising zyxin N-term and LDO constructs are indicated. **b,** Maximum intensity projections (MIPs) of confocal z-stacks of stretched zyxin null MEFs expressing the indicated eGFP-labeled constructs. Top, eGFP. Middle, F-actin (phalloidin). Bottom, endogenous VASP (immunostaining). Blue boxes highlight constructs showing visible recruitment of VASP to SFs. Double-headed arrow indicates the uniaxial stretch direction. Scale bar, 10 μm. **c,** Whole-cell actin enrichment of endogenous VASP in stretched zyxin null MEFs expressing the constructs shown in **b**. Bars represent means. 7 ≤ n ≤ 39. Dunnett’s T3 multiple comparison test after Welch’s ANOVA: NS, p > 0.05; ** p < 0.01; *** p < 0.001; **** p < 0.0001. **d,** Top, spinning-disk confocal snapshot of a zyxin null MEF co-expressing F-tractin-mApple (magenta) and zyxin-FHL3-eGFP (green). Scale bar, 10 μm. Bottom, zoomed-in view of boxed region. Arrow heads highlight SF strain sites. Scale bar, 3 μm. **e,** Time-lapse montages of SF strain sites labelled by the indicated eGFP fusion constructs. Scale bar, 3 μm. **f,** SF strain site appearance rate. Bars represent means. Welch’s ANOVA test: NS, p > 0.05. **g,** SF strain site break frequency. Gray bars represent number of repairs, while colored bars represent number of breaks. The numbers of SFFSs in each category are indicated. Fisher’s exact test: *** p < 0.001; **** p < 0.0001.

As anticipated, overexpressed zyxin LDO, FHL2, and FHL3 in zyxin null cells failed to recruit VASP (Fig. 2b, mean of localization ratio ≈ 1) in cell-stretching experiments due to the lack of VASP-interacting domains, and the Zyxin N-terminus alone failed to mechanoaccumulate on SFs due to the absence of LIM domains (Fig. 2b). Both zyxin-FHL chimeras, however, rescued VASP recruitment to a degree comparable with full-length zyxin (Fig. 2c). We next investigated the actin repair capacity of these constructs at SF strain sites (Fig. 2d, e), whose rate of spontaneous appearance was unaffected by the overexpression of any construct (Fig. 2f). Cells expressing zyxin LDO, FHL2, and FHL3 all featured short-lived strain sites (Fig. 2e, left, Supplementary Video 1) which tended to resolve in catastrophic SF breaks (Fig. 2g). The zyxin-FHL constructs, however, localized to long-lived strain sites that displayed actin recovery (Fig. 2e, right, Supplementary Video 1). Consistently, strain site break frequency was significantly reduced only in zyxin null cells expressing zyxin and zyxin-FHL chimeras (Fig. 2g). These findings collectively demonstrate that LIM domains are functionally-swappable modules which confer mechanosensitive actin localization.

### A conserved phenylalanine in each LIM domain is necessary for mechanoaccumulation

To identify features at the primary sequence level that could confer mechanoaccumulation activity, we performed multiple sequence alignments on the isolated LIM domains from the mechanoresponsive and non-mechanoresponsive LIM proteins identified in our screen. This revealed a single phenylalanine residue that is invariant in mechanoresponder LIM domains with the exception of the third LIM domain of zyxin family members, but variable in the non-mechanoresponders, suggesting it could be necessary but not sufficient for conferring mechanoaccumulation activity (Fig. 3a, Extended Data Fig. 6a). We superimposed the structures of mechanoresponder LIM2 from FHL2 and non-mechanoresponder LIM4 from LIMS1 to examine the chemical environment of residues at this conserved position. The phenylalanine (F131, green) in FHL2 LIM2 is simultaneously surface-exposed, where it would be available to mediate protein-protein interactions, and engaged in Van der Waals interactions in the small hydrophobic core of the domain, a hybrid pose also adopted by a histidine (H238, red) at the same location in LIMS1 LIM4 (Fig. 3b).

**Fig. 3.**
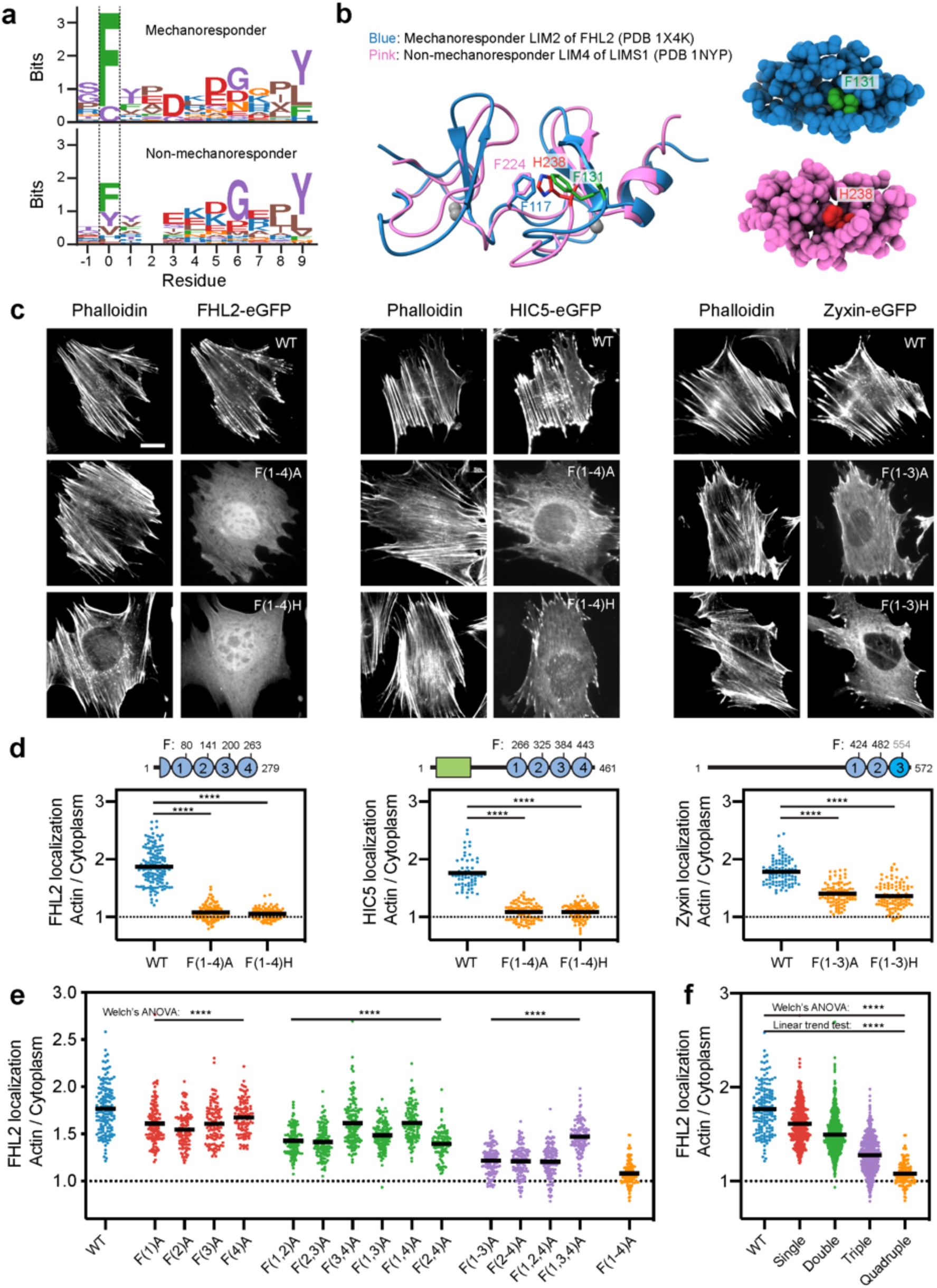
**Conserved phenylalanine residues in mechanoresponder LIM domains additively contribute to mechanoaccumulation. a**, Multiple sequence alignment logos for amino acids around the conserved phenylalanine (position 0) in mechanoresponder LIM domains (top) and the corresponding region in non-mechanoresponder LIM domains (bottom). **b**, Superimposed ribbon diagrams (left) and space-filling surface representations (right) of FHL2 LIM2 and LIMS1 LIM4. The conserved phenylalanine in FHL2 LIM2 (F131) and the histidine at the same location in LIMS1 LIM4 (H238) are highlighted in green and red, respectively. **c**, Epifluorescence micrographs of MEFs expressing the indicated eGFP-labeled FHL2 (left panel), HIC5 (middle panel), and zyxin (right panel), stained with phalloidin to label F-actin. Scale bar, 20 μm. **d**, Top: Primary sequence cartoons showing positions of conserved phenylalanines. The divergent third LIM domain of zyxin is highlighted in a different shade of blue. Bottom: Whole-cell actin enrichment of constructs shown in **c**. Left, FHL2 (84 ≤ n ≤ 151); middle, HIC5 (60 ≤ n ≤ 87); right, zyxin (90 ≤ n ≤ 101). N = 2 biological replicates. Games-Howell’s multiple comparison test after Welch’s ANOVA: **** p < 0.0001. **e**, Whole-cell actin enrichments of wild-type (blue, n = 151) and FHL2 with single (red, 112 ≤ n ≤ 138), double (green, 93 ≤ n ≤ 138), triple (purple, 109 ≤ n ≤ 121), and quadruple (orange, n = 112) phenylalanine mutations in the indicated combinations. N = 2 biological replicates. Welch’s ANOVA test: **** p < 0.0001. **f**, Actin enrichments of mutant constructs from **e**, pooled by number of lesions. Welch’s ANOVA test: **** p < 0.0001. Linear trend test^57^: **** p < 0.0001.

To investigate the functional role of these conserved phenylalanines in mechanoresponsive proteins, we generated point mutants where we substituted them with alanine or histidine (mimicking non-mechanoresponder LIMS1 LIM4) in every LIM domain. For brevity, we indicate the LIM domains featuring mutations in parenthesis, e.g., FHL2 with four phenylalanine to alanine substitutions (FHL2 F80A F141A F200A F263A) is named FHL2 F(1-4)A. We found that F(1-4)A and F(1-4)H substitutions in both FHL2 and HIC5 abolish their actin enrichment in MEF cells plated on glass (Fig. 3c, d, mean localization ratios ≈ 1), a stiff condition which activates mechanoaccumulation. Zyxin family members, however, have a unique LIM3 featuring two insertions and lacking the conserved phenylalanine (Extended Data Fig. 6a, cysteine in sequence logo position 0 of Fig. 3a, top). Nevertheless, zyxin LIM3 has a phenylalanine adjacent to the conserved position (Extended Data Fig. 6a), which we hypothesized could fulfill an analogous role in mechanoaccumulation. We first tested this hypothesis in the context of the zyxin LDO, finding that zyxin LDO F(1-3)A had more severe impairment of actin enrichment than zyxin LDO F(1,2)A versus wild-type zyxin LDO (Extended Data Fig. 6b, c). We next generated full-length zyxin F(1-3)A and F(1-3)H mutants, which showed reduced actin enrichment versus full-length wild-type (Fig. 3d). This suggests that the tandem LIM domains of zyxin operate by a fundamentally similar mechanism to other mechanoresponder LIM domains. Nevertheless, all the mutants of zyxin and zyxin LDO retain a greater degree of actin enrichment (Fig. 3d, Extended Data Fig. 6c, mean localization ratios > 1) than the equivalent mutants of other LIM proteins, suggesting that zyxin LIM3 may be specialized to confer stronger mechanoaccumulation activity through additional mechanisms. Together, these data show that the conserved phenylalanine in mechanoresponder LIM domains is necessary for their mechanoaccumulation activity.

### Tandem LIM domains contribute additively to mechanoaccumulation

All mechanoresponsive LIM proteins we identified contain at least three LIM domains in tandem, suggesting mechanoaccumulation could require the operation of multiple LIM domains in series. To test whether mechanoresponder LIM domains contribute additively to mechanoaccumulation, we systematically mutated the conserved phenylalanine to alanine in each LIM domain of FHL2 individually and in all possible combinations, and examined their localization in MEFs plated on glass (Fig. 3e, f).

Mutants with the same number of lesions exhibit distinct degrees of actin enrichment (Fig. 3e), suggesting that all LIM domains do not possess identical mechanoresponder activity. Notably, LIM2 emerged as the largest contributor to SF localization, as FHL2 F(2)A had the lowest actin enrichment among the single mutants and FHL2 F(1,3,4)A had the highest actin enrichment among the triple mutants (Fig. 3e). LIM4, on the other hand, appears to make the weakest contribution, as indicated by FHL2 F(4)A having the highest actin enrichment among the single mutants, as well as FHL2 F(3,4)A and FHL2 F(1,4)A among the double mutants (Fig. 3e). Nevertheless, pooling data from mutants with the same number of lesions reveals a linear trend, in which the actin enrichment decreases as the number of mutations increases (Fig. 3f). This suggests that all LIM domains in a tandem array contribute additively but not equally to mechanoaccumulation.

### Force along actin filaments is necessary and sufficient for LIM protein binding

We next sought to define the biophysical mechanism of LIM domain mechanoaccumulation. Hypothesizing that force across actin filaments is required to activate binding, we developed a novel adaptation of the gliding filament assay^32^ to monitor protein binding to filaments under load. Myosin Va (plus-end directed) and myosin VI (minus-end directed) motor proteins are immobilized on a glass coverslip, where they apply stress along actin filaments in the presence of ATP (Fig. 4a). We purified C-terminally Halo-tagged full-length FHL3, HIC5, and zyxin (Extended Data Fig. 7a, b), and labeled them with a fluorogenic Halo-reactive dye^33^. We then examined the ability of each protein to interact with F-actin in the absence and presence of force (controlled by supplying ATP) by total internal reflection fluorescence (TIRF) microscopy.

**Fig. 4.**
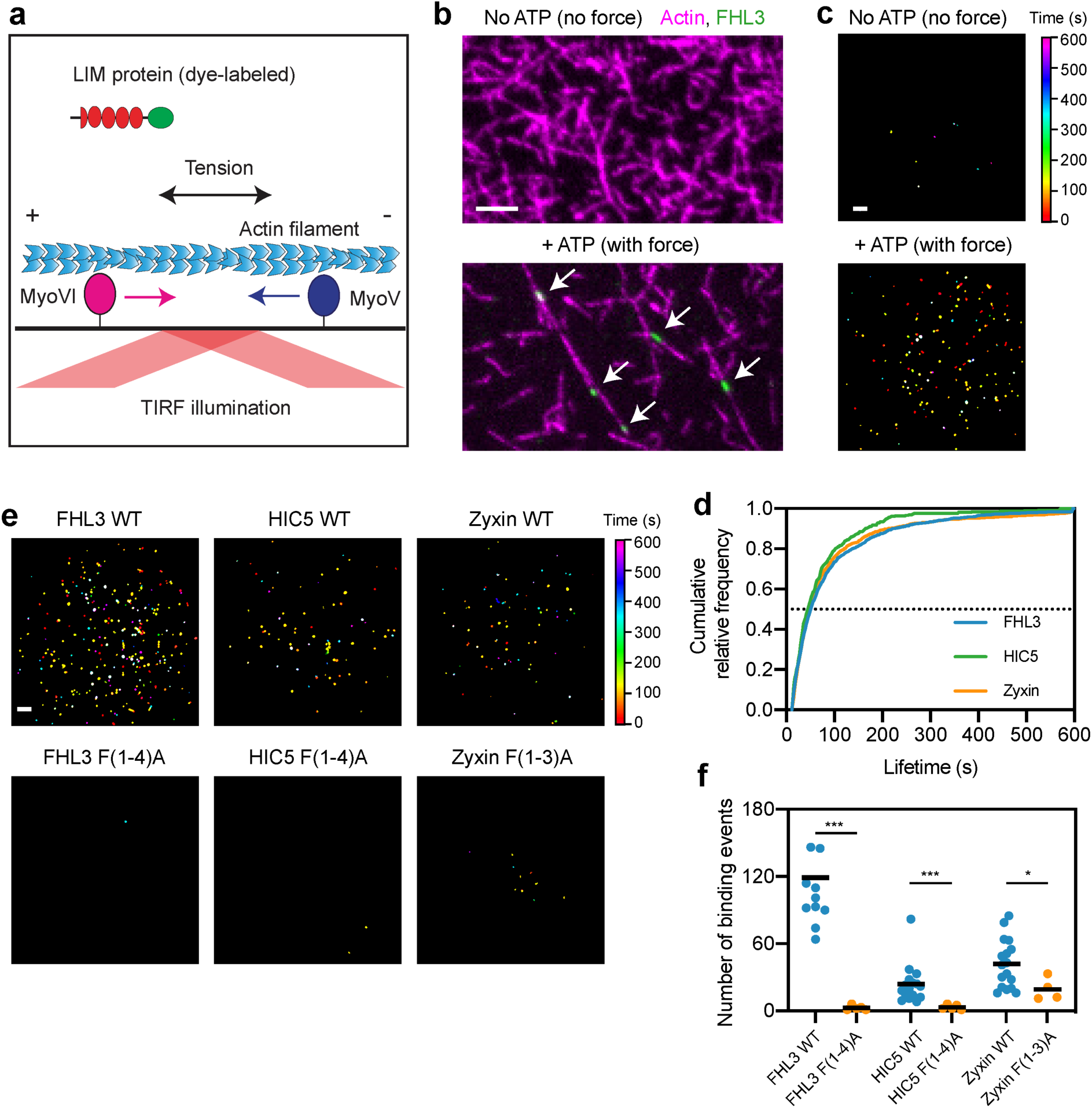
**Strain in actin filaments is necessary and sufficient for direct binding by LIM proteins. a,** Schematic of *in vitro* force reconstitution TIRF assay. **b,** TIRF snapshots of 10% rhodamine labelled actin filaments (magenta) and FHL3-Halo (green) in the absence (top) and presence (bottom) of force generation. Arrows (bottom) highlight the punctate patches of FHL3. Scale bar, 5 μm. **c,** Cumulative projections of detected patches of FHL3 in the absence (top) and presence (bottom) of 0.5 mM ATP. Scale bar, 10 μm. **d,** Cumulative relative frequency of binding lifetimes of FHL3 (blue), HIC5 (green), and zyxin (orange). Half-lives are extracted by linear interpolation; errors represent standard deviations obtained from bootstrapping (Methods). **e,** Cumulative projections of detected patches of wild-type (top) and mutant (bottom) FHL3 (left), HIC5 (middle), and zyxin (right), color-coded by time. Scale bar, 10 μm. **f,** Number of patches detected in equal imaging periods across independent trials for wild-type and mutant FHL3, HIC5, and zyxin. Bars represent means. Dunnett’s T3 multiple comparison test after Welch’s ANOVA: * p < 0.05; *** p < 0.001. One outlier in FHL3 WT is not shown.

We found that FHL3, HIC5, and zyxin all bind to F-actin in small “patches” only in the presence of ATP (Fig. 4b, Supplementary Video 2), suggesting that force is necessary and sufficient to activate this direct binding. This is further demonstrated in the cumulative projections of patches detected and tracked over time (Methods), which show many more patches in the presence of force generation (Fig. 4c). FHL3, HIC5, and zyxin patches exhibit indistinguishable lifetime distributions (Fig. 4d). The nearly identical patch half-lives (48.3 ± 1.7 s for FHL3, 42.7 ± 2.9 s for HIC5, and 44.3 ± 1.5 s for zyxin) extracted from the lifetime distributions of all three mechanoresponsive LIM proteins are consistent with them forming along regions featuring a common molecular mark in strained F-actin.

To test whether the conserved phenylalanine is required for this direct binding to strained actin, we purified Halo-tagged FHL3 F(1-4)A, HIC5 F(1-4)A, and zyxin F(1-3)A (Extended Data Fig. 7a, b). All mutants eluted with similar retention volumes to their wild-type counterparts when subjected to size-exclusion chromatography, and also displayed overlapping circular dichroism spectra, suggesting multi-site phenylalanine to alanine mutations do not grossly disrupt the structure of LIM domains (Extended Data Fig. 7a, c). As illustrated in cumulative projections of patches detected over time (Fig. 4E), all the mutants were almost completely defective in patch formation (Fig. 4f). Zyxin F(1-3)A nevertheless retained a greater residual degree of strained actin binding *in vitro* than the other two mutants (Fig. 4e, f), mirroring its incomplete loss of actin enrichment in cells versus the FHL2 F(1-4)A and HIC5 F(1-4)A constructs (Fig. 3d). These findings, coupled with the concordant properties of phenylalanine mutants in cells, suggest that direct force activated F-actin binding is the conserved mechanism of LIM domain mechanoaccumulation *in vitro* and *in vivo*.

### LIM proteins recognize a pre-break state of F-actin which spreads along filaments

To gain further insight into the mechanism of strained F-actin recognition by LIM domains, we examined the dynamics of individual patches in detail. Patches formed by FHL3, HIC5, and zyxin can initiate both in the middle (Fig. 5a, top) and at the ends of actin filaments (Fig. 5a, bottom), suggesting there is no intrinsic requirement for an exposed filament end to license patch formation. Patches appear along filaments as they straighten, consistent with the presence of mechanical tension, then disassociate abruptly upon load release and filament relaxation (Supplementary Video 3), or upon filament breakage (Fig. 5a, Supplementary Video 4). To quantify the relationship between unbinding events and filament breakage, we calculated the log ratio of the average actin intensity in the region covered by patches before unbinding vs. immediately after (Methods). A positive log ratio is consistent with LIM proteins dissociating from a filament when it breaks at the patch site, as illustrated by fluorescence intensity line scans of a representative FHL3 patch (Fig. 5c). Indeed, the vast majority of the patches formed by all three LIM proteins unbind upon filament breakage (Fig. 5d), suggesting this is the major process determining patch lifetime. The residual fraction with negative log ratios presumably results from the unbinding of LIM proteins due to filament relaxation (Supplementary Video 3).

**Fig. 5.**
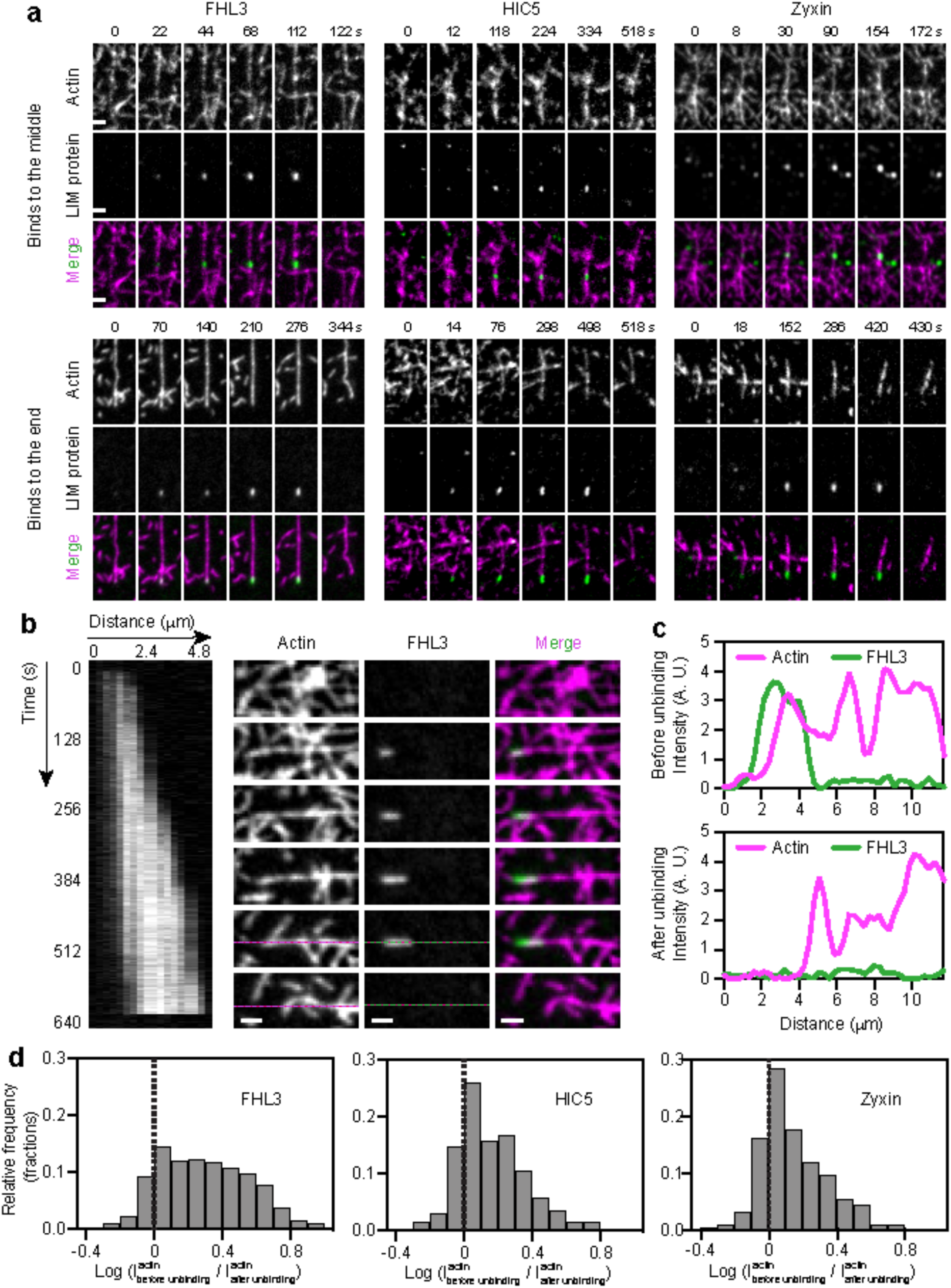
**LIM proteins bind along local filament regions featuring a pre-break state of F-actin. a,** Montages of FHL3 (left), HIC5 (middle) and zyxin (right) patches forming in the middle (top) and at the ends (bottom) of actin filaments in the presence of force generation. Scale bars, 2 μm. **b,** Kymograph (left, FHL3 channel) and montage (right) of an FHL3 patch. Displayed snapshots are at the time points indicated along the kymograph. Scale bar, 2 μm. **c,** Intensity line scan along dotted lines shown in **b** before (top) and after (bottom) the disappearance of FHL3. **d,** Histograms of the log ratio of actin intensity at detected patches before versus immediately after the disappearance of FHL3 (left), HIC5 (middle), and zyxin (right). Dash lines indicates equal actin intensity before and after LIM-protein disappearance.

It is also clear that patches are not the result of single-molecule binding events, as they are frequently anisotropic (Fig. 5b). Kymograph analysis of a representative FHL3 patch shows steady elongation at a rate of 0.3 μm / minute, followed by an absence of growth for several seconds before it dissociates in a single frame upon filament breakage (Fig. 5b, Supplementary Video 5). Occasionally, after ruptures under a patch in the middle of a filament, we observe retention of a lower level of LIM protein at one end of the two new filament segments, which could then continue to grow as load developed (Extended Data Fig. 8). We observed no apparent correlation between lifetime and maximum patch area or maximum patch intensity (Extended Data Fig. 9), indicating that in the absence of other cytosolic binding partners LIM proteins themselves do not reinforce strained actin filaments, and that patch lifetime is presumably determined by a filament’s mechanical endurance against breakage rather than the number of LIM proteins present in a patch. These observations are consistent with patches forming along local regions in filaments that feature a distinct, force-induced F-actin conformation that spreads along filaments and is recognized and bound by mechanoresponder LIM domains prior to filament rupture induced by myosin-generated forces.

### Tensed actin binding retains FHL2 in the cytoplasm in stiff environments, restricting its nuclear shuttling

Our finding that mechanoresponsive LIM proteins directly associate with a “pre-break” state of F-actin dovetails with the known actin repair function of zyxin. However, our observation that wild-type FHL protein overexpression does not rescue the repair deficiency in cells lacking zyxin (Fig. 2c) suggests that strained-actin binding plays a different role in this family. As increasing ECM stiffness is known to upregulate SF-mediated contractility^34^, we hypothesized that a concomitant increase in the cytoplasmic pool of tensed actin binding sites could act as a “sink” to retain FHL2 in the cytoplasm and suppress its nuclear translocation^16^. To test this hypothesis, we expressed eGFP-labeled wild-type FHL2 and FHL2 F(1-4)A in MEFs, and plated the cells on glass, 50-kPa, and 12-kPa hydrogels (Fig. 6a). We found that wild-type FHL2 displayed enhanced nuclear enrichment with decreasing substrate stiffness (Fig. 6a, b), consistent with the reported behavior of endogenous FHL2 in Human Foreskin Fibroblasts (HFFs)^16^. In contrast, FHL2 F(1-4)A was predominantly nuclear regardless of substrate stiffness (Fig. 6a, b). Moreover, wild-type FHL2’s actin enrichment showed an inverse trend, diminishing with decreasing stiffness as the model would predict, while FHL2 F(1-4)A always displayed low actin enrichment (Fig. 6b).

**Fig. 6.**
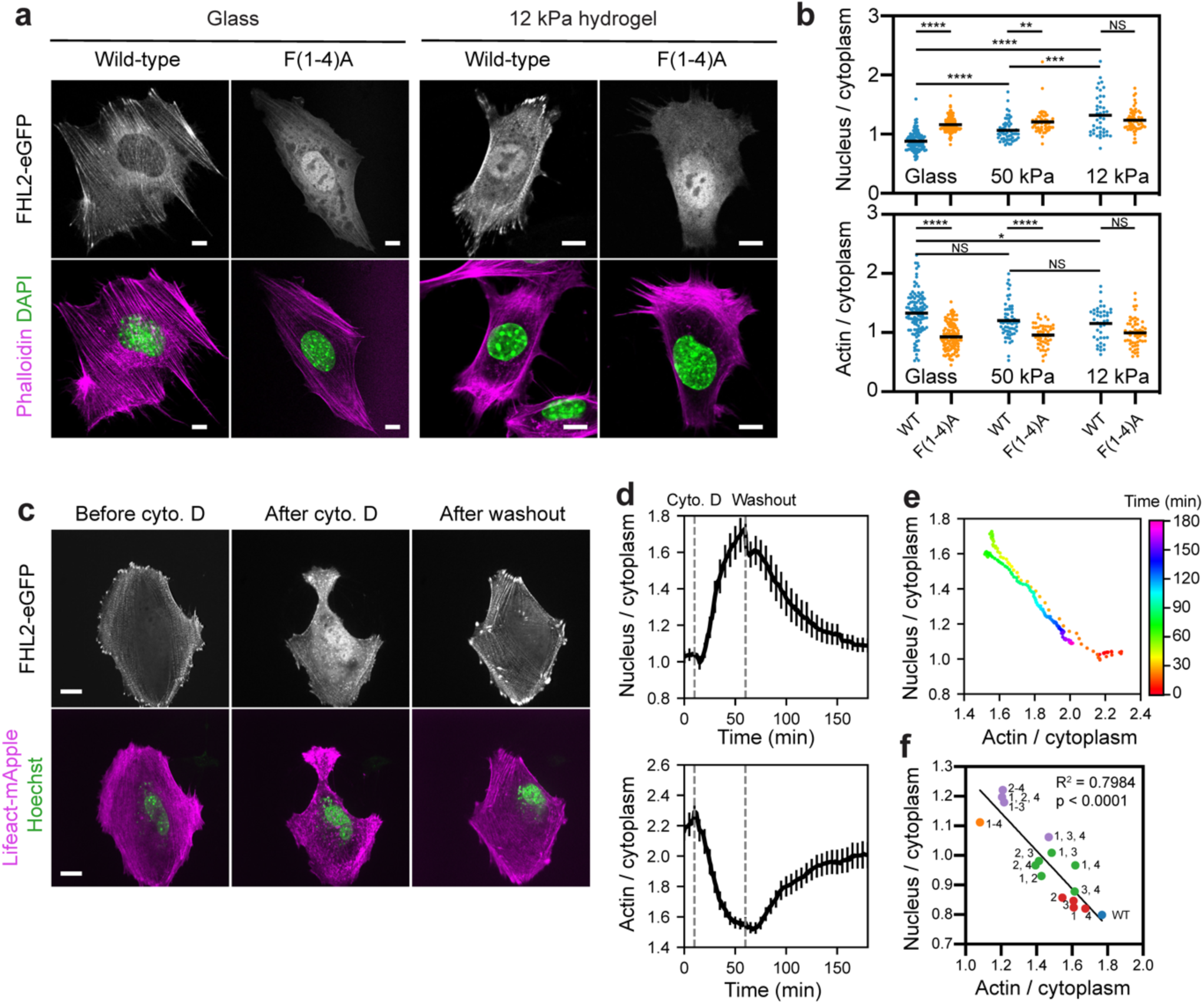
**Tensed actin acts as a sink to retain FHL2 in the cytoplasm. a**, Confocal slices of MEFs expressing eGFP-labeled wild-type and FHL2 F(1-4)A plated on substrates of varying stiffness as indicated and stained with phalloidin (magenta) to label F-actin and DAPI (green) to label nuclei. Scale bars, 10 μm. **b**, Quantification of nuclear (top) and actin (bottom) enrichment of wild-type and FHL2 F(1-4)A in MEFs (41 ≤ n ≤ 110) plated on substrates of varying stiffness. N = 2 biological replicates; bars represent means. Games-Howell’s multiple comparison test after Welch’s ANOVA: NS, p > 0.05; * p < 0.05; ** p < 0.01; *** p < 0.001; **** p < 0.0001. **c**, Spinning-disk confocal snapshots of expressed FHL2-eGFP localization in a MEF before (left) and after (middle) cytochalasin D treatment, and after washout (right). Scale bars, 20 μm. **d**, Average nuclear (top) and actin (bottom) enrichments obtained from n = 19 cells from N = 13 biological replicates as a function of time, aligned by the timepoint of cytochalasin D addition (time 10 min). Error bar represents SEM. Dash lines indicate the time points of cytochalasin D addition and washout. **e**, Replotting of average nuclear enrichment versus average actin enrichment, displayed as a scatterplot where points are color-coded by time. **f**, Scatter plot of average nuclear enrichment versus average actin enrichment of wild-type FHL2 and FHL2 with single (red), double (green), triple (purple), and quadruple (orange) phenylalanine mutations in LIM domains indicated by numbers. Data were extracted from the same set of experiments displayed in Fig. 3e. Trend line from a linear fit (R^2^ = 0.7984, p < 0.0001) is displayed.

To test if the cytoplasmic tensed actin sink model could quantitatively explain mechanosensitive FHL2 nuclear-cytoplasmic partitioning, we performed live-cell imaging of Hoechst-treated MEFs co-expressing FHL2-eGFP and LifeAct-mApple while reversibly disrupting the actin cytoskeleton with low doses of cytochalasin D. Cytochalasin D treatment triggered rapid FHL2 dissociation from the actin cytoskeleton and nuclear shuttling, which was reversed upon washout (Fig. 6c-e, Supplementary Video 6). Nuclear and actin enrichments tracked on a frame-by-frame basis show opposite trends throughout the time-course (Fig. 6d). A time-coded scatter plot of average nuclear enrichment versus average actin enrichment reveals two trajectories with opposing directionality that essentially overlap (Fig. 6e), suggesting that FHL2 nuclear localization is dictated by the number of available tensed actin binding sites. We next examined the nuclear enrichment of our allelic series of phenylalanine mutants with graded disruption of actin enrichment (Fig. 3e, f), and found a negative linear relationship between nuclear enrichment and actin enrichment (Fig. 6f). This finding suggests that FHL2 nuclear localization is also dictated by its affinity for tensed actin.

Phosphorylation of FHL2 Y93 by FAK has previously been reported to be required for FHL2 nuclear translocation in soft environments in HFFs^16^; a counterintuitive finding, as tensile force and stiff ECM collectively upregulate FAK activity^35–37^. To dissect the interplay of tensed actin binding and FAK phosphorylation, we generated phospho-resistant FHL2 mutants and pharmacologically inhibited FAK, which did not impact mechanosensitive FHL2 localization in MEFs (Extended Data Figure 10), demonstrating that strained actin binding is the dominant upstream process mediating FHL2 mechanotransduction in these cells.

## Discussion

We find that mechanoresponsive LIM proteins directly bind tensed F-actin through a conserved mechanism. This interaction excludes FHL2 from the nucleus in stiff environments (Fig. 7), providing, to our knowledge, the first direct link between upstream forces transmitted through the cytoskeleton and downstream nuclear localization of a transcriptional co-activator. Tissue stiffening is associated with tumorigenesis^38^ and cancer progression^39^, suggesting that aberrant cytoskeletal mechanotransduction through FHL2 may contribute to FHL2’s role in this disease. We report a strategy for generating point mutants specifically deficient in tensed F-actin binding that is applicable to all mechanoresponsive LIM-protein families we identified. This will facilitate the context-specific dissection of mechanoregulated gene expression controlled by FHL2, as well as by other transcriptional co-activators in the FHL family^40, 41^. Dynamic nuclear localization of members of the zyxin and paxillin families has also been reported^19, 42^, the physiological relevance of which is not clear. We speculate that their nuclear localization is coordinately regulated by their LIM domains’ intrinsic activity to bind tensed F-actin, as well as by tissue and cell-type specific mechanisms which operate through their non-LIM sequence elements. This coordinated regulation may be a general mechanism for the evolution of specialized signaling networks that incorporate LIM domains as actin force-sensing modules.

**Fig. 7.**
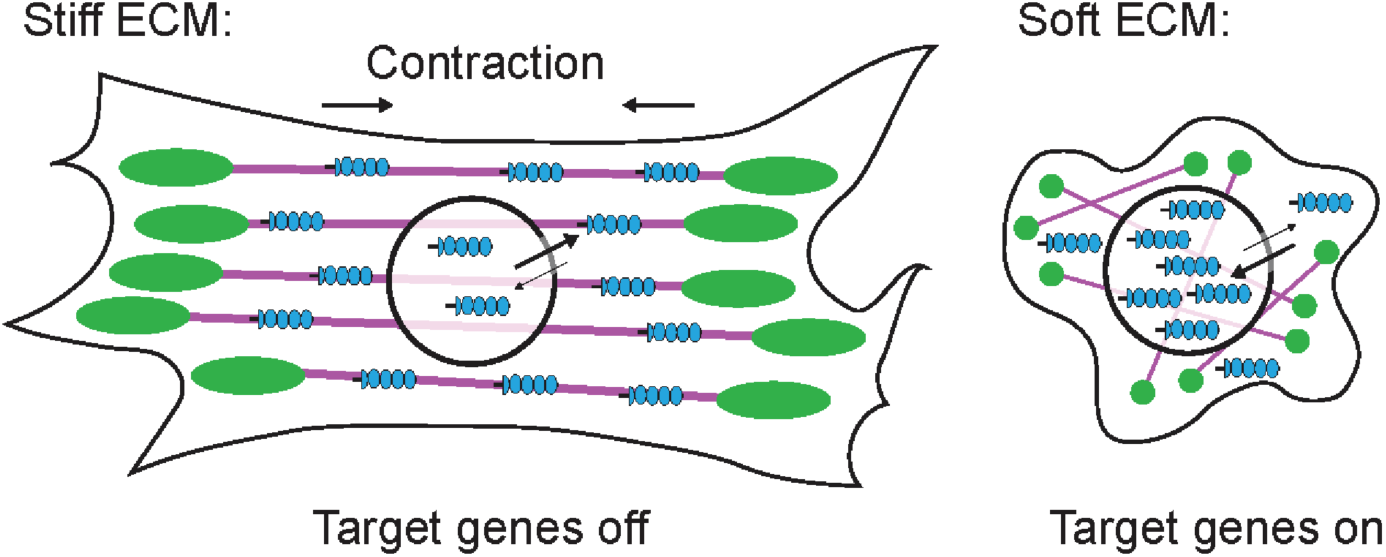
**Cartoon model of rigidity-dependent nuclear localization of FHL2.** In stiff microenvironments, FHL2 directly binds to tensed F-actin in SFs, retaining the protein in the cytoplasm. In soft microenvironments, the protein is released from SFs, where it is able to enter the nucleus and access its transcriptional targets. Magenta, SFs. Green, FAs. Blue, FHL2.

We propose that mechanoresponder LIM domains constitute a novel class of actin-binding proteins which directly bind F-actin only in the presence of mechanical force. While it is likely that additional mechanisms reinforce LIM protein mechanoaccumulation at SFs, such as high local concentrations of binding sites in F-actin bundles and interactions with other SF-localized partners, our studies establish tensed F-actin binding as the obligatory first step in this process. The most parsimonious explanation for our observation that mechanoresponsive LIM proteins bind along filaments in patches with similar kinetics is that mechanical force locally evokes a unique F-actin conformation that is specifically recognized by LIM domains. As there are dozens to hundreds of actin-binding proteins in the cell^43^, it is likely that other factors are also responsive to this conformation, providing an attractive mechanism for coordinating diverse mechanotransduction pathways through direct mechanical regulation of F-actin structure.

## Acknowledgements

We gratefully acknowledge Pinar Gurel (RU) for initial assistance with setting up the TIRF force reconstitution assays. We also thank Laura Hoffman and Chris Jensen (HCI) for the gift of the zyxin^-/-^ MEF cell line, Yasuharu Takagi and James Sellers (NHLBI) for the gift of myosin motor proteins and Luke Lavis (HHMI Janelia) for the gift of JF-646 dye for pilot TIRF studies, Laura Yen (NYSBC) for technical assistance with cloning, Matthew Reynolds (RU) for insightful discussion on statistical analysis, and Jonathan Winkelman, Ronen Zaidel-Bar, and Margaret Gardel for discussion of unpublished data. X.S. is supported by a National Institutes of Health Cancer Cell Biology training grant (CA009673-40) and a Pels Family Foundation Fellowship. R.G. is supported by a H. Li. Memorial Fellowship. L.A., A.M.P., C.M.W., and G.M.A. were supported by the National Heart Lung and Blood Institute’s Division of Intramural Research. This work was additionally supported by grants from the Irma T. Hirschl / Monique Weill-Caulier Trust and the Pew Charitable Trusts to G.M.A, and National Institutes of Health grants to M.B. (RO1GM050877) and G.M.A. (DP5OD017885).

## Author contributions

X.S. performed in vitro and mechanobiology studies, developed computational analysis tools, and analyzed all data. D.P., L.A., and S.E.R. performed cell-stretching experiments. D.P. prepared and analyzed phenylalanine mutants in cells. L.A. performed zyxin/FHL chimera experiments. M.A.S. and E.B. performed live-cell imaging studies of FHL2 nuclear localization with actin perturbation. R.G. prepared and purified LIM proteins and myosin motors for in vitro studies. R.C.C. performed live-cell imaging studies of wild-type LIM proteins. C.M.W., M.C.B. and G.M.A. contributed to study design, oversaw research, and analyzed data. G.M.A. coordinated the study. X.S. and G.M.A. wrote the paper, with input from all authors.

## Data Availability

All data supporting the findings of this study are available from the corresponding author upon reasonable request.

## Code Availability

All custom code used in data analysis is available at: https://github.com/alushinlab/LIM-domain.

## METHODS

Chemical reagents were purchased from Sigma-Aldrich unless otherwise specified.

### Molecular biology

#### Plasmids and cloning

Plasmids encoding C-terminal eGFP-labeled full-length human LIM proteins were purchased from GeneCopoeia in the m98 vector, where expression is driven by the CMV promoter. Plasmids encoding eGFP-labeled LDOs, zyxin N-term, and all phenylalanine mutants, were generated via Gibson assembly (NEB) in the same vector^44^. The coding sequences for the zyxin-FHL2 and zyxin-FHL3 were generated by gene synthesis (Blue Heron), then subcloned into the m98 vector by Gibson assembly.

For heterologous expression and purification, full-length human FHL3 (AA 1-280), HIC5 (AA 1-461), and zyxin (AA 1-572), coding sequences were subcloned into a modified pCAG mammalian expression vector with a C-terminal Halo-tag, PreScission protease cleavage site, and GFP-tag. Human myosin Va (AA 1-1091, Dharmacon BC172485) and myosin VI constructs (AA 1-1021, a generous gift from James Sellers, NHLBI) were subcloned into a modified pCAG mammalian expression vector with a C-terminal GFP-tag and Flag-tag. Human full-length calmodulin (AA 1-149, a generous gift from Roderick Mackinnon, The Rockefeller University) was subcloned into a modified pCAG mammalian expression vector with no tag.

All plasmids were propagated in NEB 5-alpha competent *E. coli* cells (NEB, C2987U), and purified by QIAprep spin miniprep kit (QIAGEN, 27106) or PureYield plasmid maxiprep system (Promega, A2392) before transfection.

### Cell maintenance and mechanobiology

#### Cell culture and transient transfection

MEFs were cultured in DMEM (Gibco, 11995-065) supplemented with 4.5 g/L D-glucose and L-glutamine, 110 mg/L sodium pyruvate, 10% fetal bovine serum (FBS), and 1% antibiotic-antimycotic (ThermoFisher, 15240062), and grown on polystyrene tissue culture dishes (ThermoFisher, FB0875712, diameter 10 cm) at 37 °C in 5% CO_2_. MEFs were harvested at 80% confluency for transient transfection. MEFs used in the cytochalasin D treatment (Fig. 6c-e) were transfected with lipofectamine. Briefly, 2 μg of FHL2-eGFP plasmid and 2 μg of LifeAct-mApple plasmid were diluted in 250 μL of Opti-MEM I Reduced Serum Medium without serum. 10 μL of lipofectamine 2000 were diluted in 250 μL of Opti-MEM I Reduced Serum Medium without serum. The diluted plasmids and diluted lipofectamine were mixed gently, incubated at room temperature for 20 minutes, and then mixed with the cells. MEFs used in other studies were transfected with 7-10 μg of LIM-protein plasmid using a Lonza 4D-Nucleofector (SE solution, program CM-137). Each plate of MEFs (2.0 × 10^6^ cells) were used for two transfection reactions. U2OS cells were cultured in McCoy’s 5A medium supplemented with 10% FBS and 1% Penicillin/Streptomycin, and grown on polystyrene tissue culture dishes (diameter 10 cm) at 37 °C in 5% CO_2_. Cells were harvested at 70% confluency for transfection. U2OS cells used in the LIM-protein SF strain site kinetic studies (Extended Data Fig. 5) were transfected with 1-2 μg of LIM-protein plasmids.

#### Cell stretching

For the initial screen (Extended Data Figs. 2, 3), MEFs transfected with LIM-protein plasmids were plated in 4-well Strex chambers (STB-CH-4W, 2 cm^2^ × 4 chambers) coated with 50 μg/mL fibronectin (Millipore, FC010), and allowed to recover overnight. A Strex STB-140-10 cell stretching system was used to apply cyclical uniaxial stretch (program 14, 10% extension, 0.5 Hz) in an incubator (37 °C, 5% CO_2_) for 2 hours. For individual SF analysis (Fig. 1 and Extended Data Fig. 5), MEFs transfected with mechanoresponder LIM-protein plasmids were allowed to recover on polystyrene tissue culture dishes overnight, then re-plated on Flexcell UniFlex culture plates coated with ProNectin (Flexcell UF-4001P) supplemented with 10 μg/mL fibronectin, allowed to recover for 2 hours, then subjected to cyclical uniaxial stretch on a Flexcell FX6000-T apparatus (Program FX-5000, 10% extension, 0.5 Hz) in an incubator (37 °C, 5% CO_2_) for 1 hour.

#### Cell fixation and staining for fluorescent imaging

Cells were fixed in Cytoskeleton Buffer (100 mM MES pH 6.1, 1.38 M KCl, 30 mM MgCl_2_, 20 mM EGTA) with the indicated supplements. All the permeabilizing and staining steps were performed in DPBS (Gibco, 14190-144). Both unstretched and stretched MEFs were immediately fixed using 4% formaldehyde (ThermoFisher, 28908) for 20 minutes, permeabilized in 0.1% Triton X-100 for 10 minutes, incubated in 0.1 M glycine for 10 minutes, and washed in DPBS. Cells were than blocked in 2% Bovine serum albumin (BSA, Gemini Bio-Products, 700-101P) for 1 hour, stained with 165 nM Alexa Fluor 568 phalloidin (ThermoFisher, A12380) for 20 minutes and 300 nM DAPI (ThermoFisher, D1306) for 15 minutes, and washed with DPBS. The membrane of the stretching chamber was excised and mounted onto a No. 1.5 glass coverslip using DAKO mounting medium (Agilent, S3023) for imaging.

The endogenous FHL2 in MEFs was stained with anti-FHL2 monoclonal antibody (1:100, rabbit host, Abcam, ab202584) at 4 °C overnight, and Alexa Fluor 488 goat anti-rabbit secondary antibody (ThermoFisher, A32731) at room temperature for 1 hour, followed by staining with Alexa Fluor 568 phalloidin and DAPI, and mounting in DAKO mounting medium as described above.

For experiments that involve plating cells on glass or hydrogel substrates (Matrigen), the substrates were coated with 10 μg/mL fibronectin. Cells were plated on those substrates after transfection and were allowed to recover overnight. Cells were fixed and stained using the approach described above, and were then transferred to DPBS for imaging.

#### Pharmacological treatments

For live-cell imaging experiments visualizing nuclear localization dynamics of FHL2 in the presence of actin disruption (Fig. 6c-e), MEFs co-expressing FHL2-eGFP and Lifeact-mApple were incubated with 5 μg/mL Hoechst (Invitrogen, H1399) for 1 hour, then treated with 250 nM cytochalasin D (Millipore, 504776).

FAK activity of MEFs (Extended Data Fig. 10b, c) was inhibited by incubating the cells with 1 μM PF573228 (Sigma-Aldrich, PZ0117) for 1 hours. Cells were then fixed and stained for confocal imaging.

### Cellular imaging

#### Fixed cell imaging

MEFs used in the initial screen (Extended Data Figs. 2, 3) and combinatorial mutation studies (Fig. 3e, f) were imaged via epifluorescence on a Nikon Ti-E microscope using a CFI Apo 100× TIRF oil immersion objective (NA 1.49). Epifluorescence illumination was provided by a PEKA LED illuminator (Lumencor, 300 mW). Images were captured at a depth of 12 bit using a Zyla 4.2 sCMOS camera (Andor). Image acquisition was performed using the NIS-Elements software (Nikon). A z-stack that spaned 1.4-1.8 μm with a step size of 0.2 μm was acquired for each cell to ensure that the epifluorescence image with the maximum amount of actin cytoskeleton in focus (visible as maximum sharpness in the actin channel) was captured for the subsequent image analysis. Excitation power and exposure time were maintained constant to ensure fair comparisons between unstretched and stretched cells.

MEFs used in the individual SF analysis (Fig. 1 and Extended Data Fig. 5) and zyxin-FHL chimera studies (Fig. 2), were imaged on Spinning Disk System 1: a Nikon TE2000E2 microscope equipped with a spinning-disk confocal scan head (Yokogawa) using an Apo 60× TIRF oil immersion objective (NA 1.49). Illumination was provided by solid-state 488-nm (100 mW) and 561-nm (550 mW) lasers. Images were captured at a depth of 16 bit using an Myo CMOS camera (Photometrics). Image acquisition was performed using the MetaMorph software (MDS Analytical Technologies).

MEFs used in the substrate stiffness (Fig. 6a, b), FAK-phospho-resistant mutant (Y93F) (Extended Data Fig. 10a), and FAK inhibition (Extended Data Fig. 10b, c) studies were imaged on a Leica TCS SP8 confocal microscope using a Plan Apo 40× water immersion objective (NA 1.10) with motorized correction collar. Illumination was provided by a UV diode laser (405 nm, 50 mW) and a fully-tunable white-light laser (470-670 nm, 1.5 mW) with an acousto-optical modulator. Images were captured at a depth of 16 bit using HyD and PMT detectors. A z-stack was acquired for each cell with the system-optimized step size. Each image was sampled at the Nyquist frequency. Image acquisition was performed using the LAS-AF software (Leica).

#### Live cell imaging

MEFs used in zyxin-FHL chimera studies (Fig. 2) as well as U2OS cells used in the LIM-protein binding kinetics studies (Extended Data Fig. 4) were imaged on Spinning Disk System 1 as described above. Time-lapse confocal images were captured at an interval of 10 s with the cells incubated in DMEM (10% FBS, 25 mM HEPES pH 7.2, 30 U/mL oxyrase) for MEFs or McCoy’s 5a (10% FBS, 25 mM HEPES pH 7.2, 30 U/mL oxyrase) for U2OS cells without phenol red at 37 °C.

MEFs used for visualizing nuclear localization dynamics of FHL2 in the presence of actin disruption (Fig. 6C) were imaged on an Andor spinning disk confocal on a Nikon TiE microscope using a Plan Apo 60× oil immersion objective (NA 1.4). Illumination was provided by 405-nm, 488-nm, and 568-nm lasers, switched by an acousto-optic tunable filter-based laser combiner (Andor), and delivered to the spinning-disk confocal scan head (CSU-10, Yokogawa). Images were captured at a depth of 16 bit every 60 s using an iXon885 EMCCD camera (Andor). Image acquisition was performed using the Andor IQ imaging software (Andor). Cells were incubated in DMEM/F12 (15 mM HEPES) without phenol red at 37 °C. 250 nM cytochalasin D was added at 10 minutes and washed out 60 minutes after the time-lapse imaging started. Cells were imaged for another 2 hours following washout.

### *In vitro* protein studies

#### Protein expression and purification

FreeStyle 293-F cells (ThermoFisher, R79007) were cultured in FreeStyle 293 expression medium (ThermoFisher, 12338026) on an orbital shaker at 37 °C in 8% CO_2_. The cells were transfected when cell density reached 1.8×10^6^ cells/mL. For 400 mL of cell culture, 400 μg of plasmid was premixed with 1.2 mL of 1 mg/mL PEI MAX (Polysciences) in 15 mL of FreeStyle 293 expression medium, and was incubated at room temperature for 20 minutes before transfection. Myosin Va and myosin VI were co-transfected with calmodulin at a mass ratio of 1:6. Cells were grown on an orbital shaker at 37 °C in 8% CO_2_ and were harvested 60 hours after transfection.

All subsequent steps of protein purification were conducted at 4 °C or on ice unless otherwise specified. To purify Halo-tagged wild-type and mutant FHL2, HIC5, and zyxin, cells were resuspended in Lysis Buffer A (50 mM Tris-HCl pH 8.0, 150 mM NaCl, 0.2% NP40, 3 mM DTT, 2 mM ATP, 1 mM PMSF, 1 μg/mL aprotinin, leupeptin, and pepstatin), and were incubated on a rocker for 40 minutes. Cell lysate was clarified by centrifugation (Sorvall) at 20,000 g for 30 minutes. The supernatant was mixed with NHS-activated Sepharose 4 Fast Flow resin (GE Healthcare) coupled to a GFP nanobody expressed and purified by published protocols^45^ and incubated on a rocker for 1.5 hours. The resin was then washed three times with Lysis Buffer A, and incubated with 0.5 mg/mL PreScission protease that was expressed and purified by published protocols^46^ on a rocker overnight to cleave the C-terminal GFP tag. Protein was then eluted with Elution Buffer (50 mM Tris-HCl pH 8.0, 150 mM NaCl, 0.2% NP40, and 3 mM DTT). The eluted protein was sequentially purified by a Mono Q anion exchange column (GE Healthcare) followed by size exclusion chromatography on a Superdex 200 increase column (GE Healthcare) in Gel Filtration Buffer (10 mM Tris-HCl pH 8.0, 100 mM NaCl, and 3 mM DTT). Peak fractions were concentrated (Amicon Ultra-4, MWCO 10 kD) to 1-2 mg/mL, snap-frozen in liquid nitrogen, and stored at −80 °C until use.

To purify myosin Va and myosin VI, cells were resuspended in Lysis Buffer B (50 mM Tris-HCl pH 8.0, 150 mM NaCl, 2 mM MgCl_2_, 0.2% CHAPS, 3 mM DTT, 2 mM ATP, 1 mM PMSF, 1 μg/mL aprotinin, leupeptin, and pepstatin), and were incubated on a rocker for 40 minutes. Cell lysate was clarified by centrifugation (Sorvall) at 20,000 g for 30 minutes. The supernatant was mixed with anti-Flag M2 affinity beads (Sigma-Aldrich) and incubated on a rocker for 1.5 hours. The protein-bound beads were washed three times with Myosin Buffer (50 mM Tris-HCl pH 8.0, 150 mM NaCl, 2 mM MgCl_2_, and 2 mM ATP). Protein was eluted with Myosin Buffer supplemented with 100 μg/mL Flag peptide (Sigma-Aldrich). The eluent was buffer-exchanged to Storage Buffer (10 mM Tris-HCl pH 8.0, 100 mM NaCl, 2 mM MgCl_2_, and 3 mM DTT) using a concentrator (Amicon Ultra-4, MWCO 50 kD). Myosin Va and myosin VI were snap-frozen in liquid nitrogen and stored at −80 °C until use.

For SDS-PAGE analysis, 3 μg of protein was loaded to each well of a NuPAGE 12% Bis-Tris gel (Invitrogen, NP0343BOX) and stained with Coomassie (Extended Data Fig. 7b). The gel was run at 140 V for 90 minutes. All protein concentrations were estimated by Bradford colorimetric assay (ThermoFisher, 1856209), calibrated with BSA.

#### Circular dichroism spectroscopy

Halo-tagged wild-type and mutant FHL3, HIC5, and zyxin were buffer-exchanged into 100 mM NaF (Alfa Aesar, A13019) dissolved in phosphate buffer (pH 7.2) using a concentrator (Amicon Ultra-4, MWCO 10 kDa), and diluted to 0.01-0.1 mg/mL using the same solution. Spectra were collected in an AVIV circular dichroism spectrometer (model 62DS) using a cuvette with an optical path length of 1 mm. Each sample was scanned twice from 180 nm to 260 nm with a step size of 0.5 nm. The average of the two measurements is plotted in Extended Data Fig. 7c.

#### Force reconstitution assay and TIRF microscopy

No. 1.5 24 × 60 mm glass coverslips (Corning, CLS2980246) were sonicated (Branson 3800) in 100% acetone for 30 minutes, 100% ethanol for 10 minutes, and 2% Hellmanex for 2 hours. They were then rinsed with MilliQ water and incubated in a solution that contains 1 mg/mL mPEG-silane (Laysan bio, MPEG-SIL-5000), 10 mM HCl, and 96% ethanol on a shaker overnight at room temperature. Individual coverslips were sequentially rinsed in ethanol and water, air-dried, and stored in nitrogen atmosphere in a sealed container at 4 °C before use.

Actin was purified from chicken skeletal muscle as previously described^47^ and was maintained in G-Ca buffer (2 mM Tris-Cl pH 8.0, 0.5 mM DTT, 0.2 M ATP, 0.01% NaN_3_, 0.1 mM CaCl_2_) at 4 °C before use. 10% rhodamine-labeled F-actin was prepared by co-polymerizing 0.9 μM unlabeled actin monomers and 0.1 μM rhodamine-labeled actin monomers (Cytoskeleton, AR05) in the presence of G-Mg (2 mM Tris-HCl pH 8.0, 0.5 mM DTT, 0.2 mM ATP, 0.01% NaN_3_, 0.1 mM MgCl_2_) and KMEI (50 mM KCl, 1 mM MgCl_2_, 1 mM EGTA, 10 mM imidazole pH 7.0) at room temperature for 1 hour. 20 μg of rhodamine-labeled actin monomer was resuspended in 20 μL of 0.9× G-Ca buffer, incubated at 4 °C for at least 1 hour, and clarified by ultracentrifugation at 100,000 rpm in a TLA100 rotor for 20 minutes before use. F-actin was freshly polymerized for each experiment.

Myosin Va-GFP and myosin VI-GFP were premixed and diluted to 0.02 μM and 0.04 μM, respectively, with Motility Buffer (MB: 20 mM MOPS pH 7.4, 5 mM MgCl_2_, 0.1 mM EGTA, 50 mM KCl, 1 mM DTT). Halo-tagged wild-type and mutant FHL3, HIC5, and zyxin were fluorescently labelled by incubation with Janelia Fluor 646 HaloTag ligand (JF646, Promega, GA1121) at a 1:2 molar ratio on ice for 2 hours. Proteins were freshly labeled for each experiment. Excess dye was removed using Pierce dye removal columns (ThermoFisher, 22858). JF646-labeled proteins were then diluted to 500 nM with Imaging Buffer (MB supplemented with 15 mM glucose, 100 μg/mL glucose oxidase, 20 μg/mL catalase, and 1 μM calmodulin). Proteins were clarified by ultracentrifugation at 60,000 rpm in a TLA100 rotor for 15 minutes at 4 °C immediately before imaging.

Imaging wells were prepared by attaching a CultureWell reusable gasket (GraceBio 103280, diameter 6 mm, well depth 1 mm) to the PEG-coated coverslip. Before imaging, each well was treated with the following sequence: coated with 0.02 μM GFP-myosin Va and 0.04 μM GFP-myosin VI in MB for 2 minutes, blocked with 0.1% polyvinylpyrrolidone in MB for 1 minute, coated with 0.8 μM 10% rhodamine-labeled F-actin in MB for 30 s, rinsed with MB, then immersed in 20 μL of imaging buffer without ATP for TIRF microscopy. Sequential solutions were added and removed by pipetting.

The imaging well was mounted on a Nikon Ti-E microscope equipped with an H-TIRF motorized module. Time-lapse dual-color TIRF imaging was then initiated to visualize basal level background, followed by addition of 20 μL of 500 nM JF646-labeled LIM protein in the presence or absence of 1 mM ATP (final concentrations: 0.5 mM ATP, 250 nM JF646-labeled LIM protein) in the “+ Force” and “– Force” conditions, respectively. This ensured that the onset of binding events was captured in the “+ Force” condition. Images were acquired every 2 s through a CFI Apo 60× TIRF oil immersion objective (NA 1.49), a quad filter (Chroma), and an iXon EMCCD camera (Andor) with the Perfect Focus System (Nikon) engaged. Illumination was provided by 561-nm (50 mW) and 640-nm (45 mW) lasers (Agilent) switched by an acousto-optic tunable filter. Image acquisition was performed using the NIS-Elements software (Nikon).

## DATA ANALYSIS

Image analysis was performed with custom Python scripts utilizing functions from the scikit-image package^48^ unless otherwise specified.

### Analysis of cell images

#### Actin enrichment

To identify mechanoresponder LIM proteins from the screen (Extended Data Figs. 2, 3), actin enrichment of the LIM protein in each cell was computed from a z-stack of epifluorescence images. The maximum intensity projection (MIP) of the eGFP channel was thresholded (li method), binarized, and subsequently applied as a cell mask to the DAPI and phalloidin channels to exclude the interference from the surrounding cells with negligible levels of eGFP-labeled LIM protein expression. To find the slice with the largest fraction of the actin cytoskeleton in focus in the z-stack, the Sobel transform was used on the phalloidin channel to find the edge magnitude (gradient of the raw image). The sharpness of each image in the phalloidin channel is defined as the sum of the edge magnitude normalized by the sum of the pixel intensity of the raw image. eGFP, phalloidin, and DAPI images obtained at the z-height that conveys the maximum sharpness in the phalloidin channel were then used to compute actin enrichment.

The actin and nuclear masks were generated by thresholding and binarizing the phalloidin (mean method) and DAPI (li method) channels, respectively. To partition the contribution of LIM proteins localized in the nucleus, a “clean” actin mask was generated by excluding the nuclear region from the actin mask. A cytosolic mask was generated by excluding the nuclear and actin region from the cell mask. The clean actin mask and cytosolic mask were individually applied to the eGFP image obtained at the same z-height. The actin enrichment was computed as the ratio of the average intensity of eGFP on actin cytoskeleton to that in the cytosol (Extended Data Fig. 1a). eGFP images with more than 500 saturated pixels covered by the cell mask were not considered for quantification. The same approach was used to quantify the actin enrichments of combinatorial point mutants (Fig. 3d-f).

Cell and actin masks were generated by thresholding and binarizing the MIP of confocal z-stacks of the eGFP (LIM protein, mean method) and phalloidin (actin, li method) channels, respectively. Cytosolic mask was generated by excluding the actin region from the cell mask. The actin enrichment of VASP (Fig. 2c) was computed using the same approach described above.

#### SF enrichment

To quantify the enrichment of LIM proteins on individual SFs (Fig. 1c-e), a polygonal ROI that encompasses the targeted SF and a sub-stack of confocal slices that contain the targeted SF were extracted manually in FIJI^49^. The nuclear region was intentionally avoided in the ROIs. A custom FIJI Macro script was used to generate ROI masks and MIPs of the eGFP (LIM protein) and phalloidin (actin) channels with background subtraction (rolling ball radius, 50 pixels) through batch processing in FIJI. The SF mask was generated by thresholding (yen method) and binarizing the MIP of the phalloidin channel within the ROI mask. The local cytosolic mask was generated by excluding the SF region from the ROI mask. The SF and local cytosolic masks were individually applied to the MIP of the eGFP channel. The SF enrichment was computed as the ratio of the average intensity of eGFP on SF to that in the local cytosol (Extended Data Fig. 1b).

#### Nuclear enrichment

To quantify the actin and nuclear enrichments of eGFP-labeled wild-type and F(1-4)A FHL2 in MEFs plated on substrates of various stiffnesses (Fig. 6b), the MIP of the eGFP channel was thresholded (li method), binarized and subsequently applied as a cell mask to the DAPI and phalloidin channels. The confocal slices of the DAPI channel were binarized using a single threshold value, which was obtained by thresholding (isodata method) a 2-dimensional array constructed by concatenating all the slices of the DAPI channel. The area of the nucleus was computed for each slice using the binarized DAPI z-stack. Slices captured at the z-height with the largest nuclear area were used to compute the actin and nuclear enrichments. The actin enrichment was obtained using the same approach described in the initial screen. To quantify the nuclear enrichment, the nuclear mask was dilated, bounded by the cell mask, until the area of the dilated mask is twice that of the original mask. A local cytosolic mask was generated by excluding the actin and nuclear region from the dilated mask. Nuclear enrichment was computed as the ratio of the average intensity of eGFP in the nucleus to that in the local cytosol. eGFP images with more than 500 saturated pixels covered by the cell mask were not considered for quantification.

#### Live-cell imaging quantification

To quantify the timing of the mechanoaccumulation of LIM proteins relative to zyxin-FusionRed (Extended Data Fig. 4c), flashes were identified in the zyxin-FusionRed channel. ROIs were manually delineated based on each frame in eGFP and zyxin-FusionRed channels. The integral intensities within each ROI throughout the lifetime of the flash were extracted for both channels using FIJI, normalized to the maximum value in the corresponding channel, and plotted against time. The differential between the timing of the peak intensity of eGFP and that of zyxin-FusionRed were extracted from the intensity plots (Extended Data Fig. 4b).

To quantify the actin and nuclear enrichment of FHL2 in the presence of actin disruption (Fig. 6c), cell, actin and nuclear masks were generated by thresholding and binarizing the FHL2-eGFP (triangle method), Lifeact-mApple (li method), and Hoechst (isodata method) channels, respectively. Cytosolic masks were generated by excluding the actin and nuclear regions from the cell masks. Actin and nuclear enrichments were computed as described above. The colored scatter plot of nuclear versus actin enrichment was constructed in matplotlib^50^.

### Analysis of in vitro TIRF images

#### Detecting and tracking LIM protein patches

Images of the LIM-protein channel were sequentially Sobel-transformed, thresholded (triangle method), and binarized. The holes of the resulting rings (regions with large edge magnitude) were filled, producing “patch masks” that potentially represent LIM protein patches (Extended Data Fig. 1c). Each candidate patch was labeled and tracked based on the overlap of pixels between patch masks in neighboring frames. Due to the low signal-to-noise ratio of the LIM-protein channel, dim patches in the raw frames tended to be lost during tracking. Therefore, overlap was searched among the previous and future 20 mask frames (*f_t-20_* to *f_t+20_*). Based on the overlap, depth first traversal was then implemented to assign masks to patches and extract the entire trajectory of each patch.

Actin masks were generated by thresholding (mean method) and binarizing the actin channel, and were applied to the corresponding patch masks. Patches featuring no pixels which overlap with the actin mask for at least one time point likely represent non-specific binding of protein aggregates to the coverslip and therefore were excluded from further analysis. Patches that persisted to the last frame were also removed, as their lifetimes could not be measured. Furthermore, any patch that persisted for less than 5 frames was considered indistinguishable from noise and was not considered in the analysis (lifetime cutoff, 10 s).

To account for the intrinsic non-uniform illumination of TIRF, the average intensities of each patch throughout its trajectory were normalized by the corresponding average intensities of its local background. Dim patches tended to fluctuate in intensity during the time-lapse, compromising the accuracy of tracking. Therefore, an intensity cutoff was also imposed on the lifetime analysis. Trajectories with a maximum normalized patch intensity of less than 1.2 were considered too dim to be tracked accurately and were excluded from further analysis (normalized intensity cutoff, 1.2).

#### Quantifying lifetime of LIM-protein patches and half-life bootstrapping

The lifetime of each patch was then computed from the number of frames in which it was detected. The reported average half-lives (Fig. 4d) were extracted by linear interpolation on the observed cumulative probability distributions (CPDs) of lifetime. Bootstrapping was performed to estimate the standard deviations of the half-lives. Briefly, the CPD of the lifetime population was estimated by the CPD of the observed lifetimes. Samples were drawn with replacement from the observed CPD. The size of each sample equals the size of the observed data set. Resampling was performed 50,000 times, resulting in 50,000 average half-lives extracted by linear interpolation. The standard deviation of the 50,000 half-lives was reported for each LIM protein (Fig. 4d).

The maximum area and maximum normalized intensity of each trajectory were extracted by the regionprops function in scikit-image package. The scatter plots of maximum area and maximum normalized intensity versus lifetime (Extended Data Fig. 8) were constructed using matplotlib.

#### Quantifying actin-intensity change upon LIM-protein patch unbinding

Patch masks were applied to the actin channel to extract the average actin intensity for each patch. The mean of all the average actin intensities throughout the entire trajectory (*f[t_0_]* to *f[t_n_]*) of the patch was defined as the actin intensity before patch unbinding (*I_before unbinding_*). To calculate the actin intensity after patch unbinding, the last patch mask was applied to the actin channel for the future 20 frames after patch unbinding, resulting in 20 average actin intensities (*f[t_n+1_]* to *f[t_n+20_]*) covered by the last patch mask. The mean of those 20 extracted average actin intensities was defined as the actin intensity after the patch unbinding (*I_after unbinding_*). The ratio of the actin intensity before to after patch unbinding was then computed. A ratio greater than 1 indicates a decrease in actin intensity upon patch unbinding. The histogram of the logarithm of the ratio (to ensure unbiased distribution) was plotted for each LIM protein (Fig. 5d).

### Sequence alignment and molecular graphics

Multiple sequence alignment was performed using Clustal Omega^51^ provided by the European Bioinformatics Institute (EMBL-EBI) web server^52^ (https://www.ebi.ac.uk/Tools/msa/). The aligned sequences (Extended Data Fig. 6a) were colored in Jalview^53^ (http://www.jalview.org/). Alignment logos (Fig. 3a) were created using the web-based application WebLogo^54^ (https://weblogo.berkeley.edu/).

The superimposed ribbon diagrams and space-filling surface representations of FHL2 LIM2 and LIMS1 LIM4 (Fig. 3b) were created using UCSF ChimeraX^55^ (http://www.rbvi.ucsf.edu/chimerax/).

### Statistical analysis

To identify mechanoresponder LIM proteins from the screen, Dunnett’s T3 multiple comparison test after Welch’s ANOVA was performed by comparing the actin enrichments of each LIM protein in unstretched versus stretched cells simultaneously for all the LIM proteins covered in the screen (Extended Data Figs. 2, 3). NS (not significant), p > 0.05; * p < 0.05; ** p < 0.01; *** p < 0.001.

In the individual SF analysis, Games-Howell’s multiple comparison test after Welch’s ANOVA was performed by comparing the actin enrichments of each LIM protein in unstretched versus stretched cells simultaneously for all the LIM proteins belonging to the same family. Given the large number of SF ROIs extracted (n_min_ = 28, n_max_ = 244), outliers were identified by ROUT method^56^ using GraphPad Prism. 0 ≤ n ≤ 14 outliers were identified in each data set: for the numbers of data points and outliers in each data set, see Extended Data Table 1. Statistical tests were performed on the cleaned data sets with outliers removed. NS (not significant), p > 0.05; ** p < 0.01; *** p < 0.001; **** p < 0.0001.

Unless otherwise noted, all statistical analysis and plotting was performed in GraphPad Prism. The figure legends contain details of other standard statistical analyses which were employed as indicated throughout the study.

---

**Extended Data Fig. 1.**
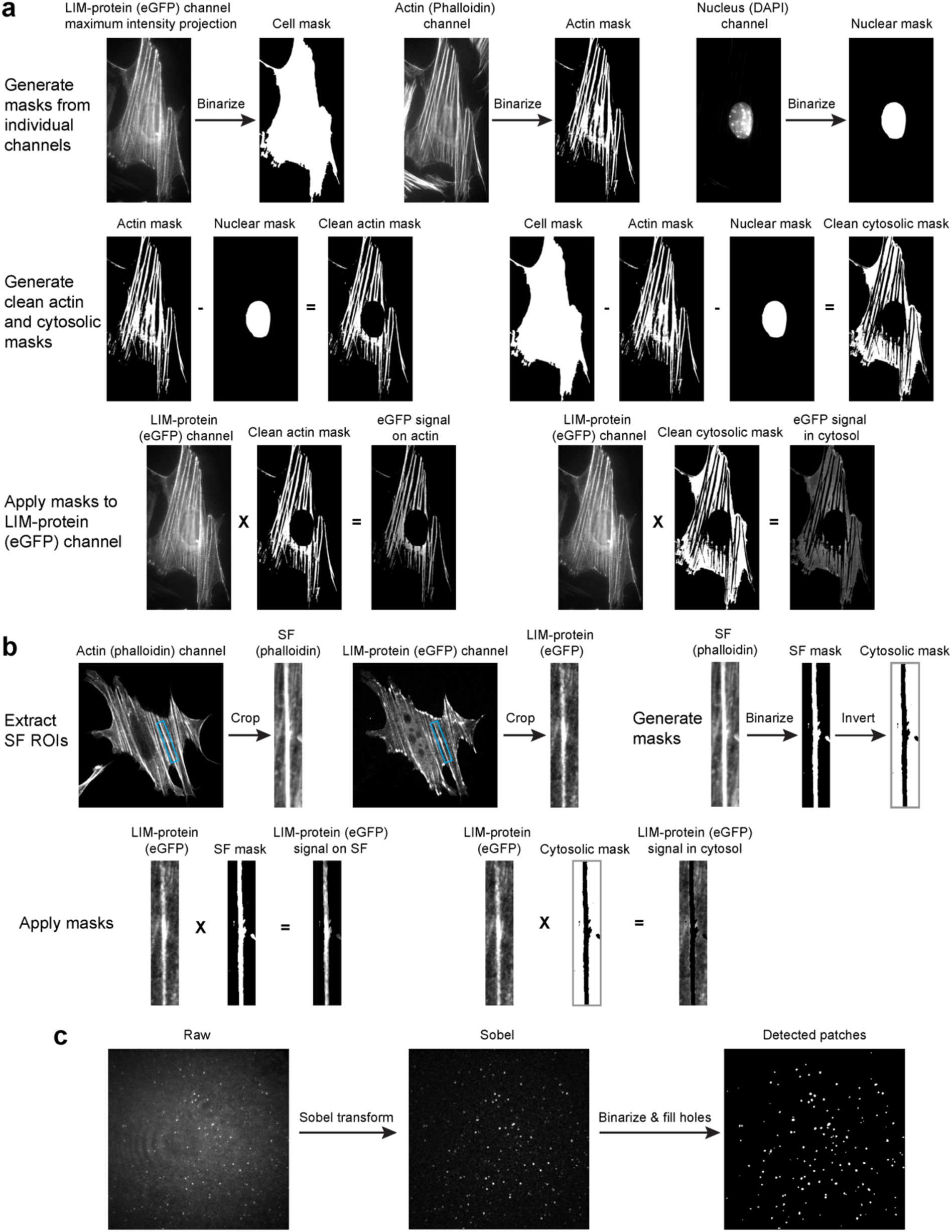
**Image analysis pipelines. a**, Single-cell-based and **b**, single-SF-based quantification of actin enrichment. **c**, Detecting LIM-protein patches in the force reconstitution assay.

**Extended Data Fig. 2.**
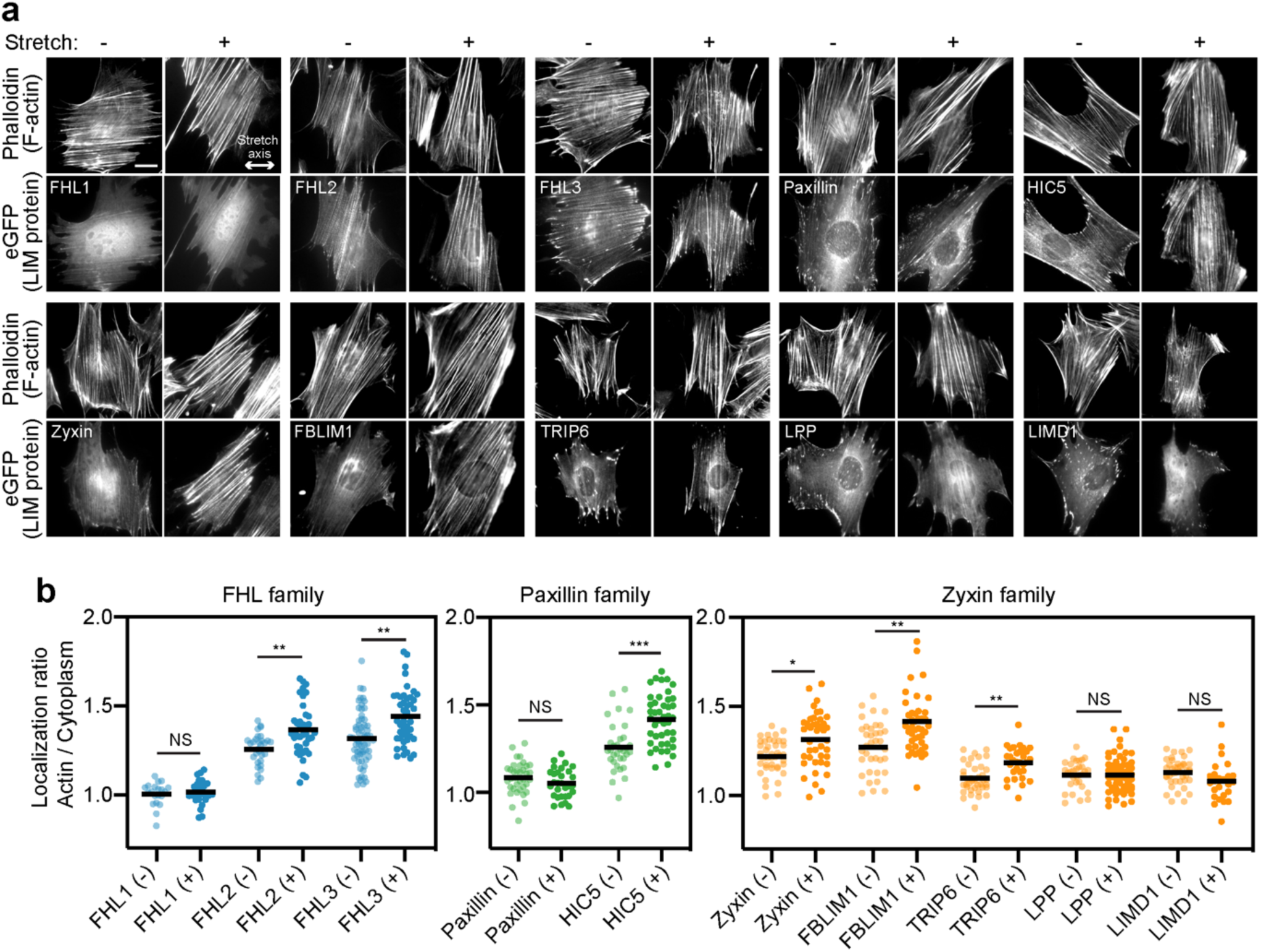
**Mechanoaccumulating LIM-protein families identified in the cell-stretching screen. a**, Epifluorescence micrographs of eGFP-labeled FHL-, paxillin-, and zyxin-family proteins in unstretched (-) and stretched (+) MEFs stained with phalloidin to label F-actin. Double-headed arrow indicates the uniaxial stretch direction. Scale bar, 20 μm. **b**, Whole-cell actin enrichment of FHL-, paxillin-, and zyxin-family proteins in unstretched (-) and stretched (+) MEFs (20 ≤ n ≤ 63). Bars represent means. Dunnett’s T3 multiple comparison test after Welch’s ANOVA is performed by comparing the actin enrichment of each LIM protein in unstretched versus stretch cells simultaneously for all LIM proteins covered in the screen. NS, p > 0.05; * p < 0.05; ** p < 0.01; *** p < 0.001.

**Extended Data Fig. 3.**
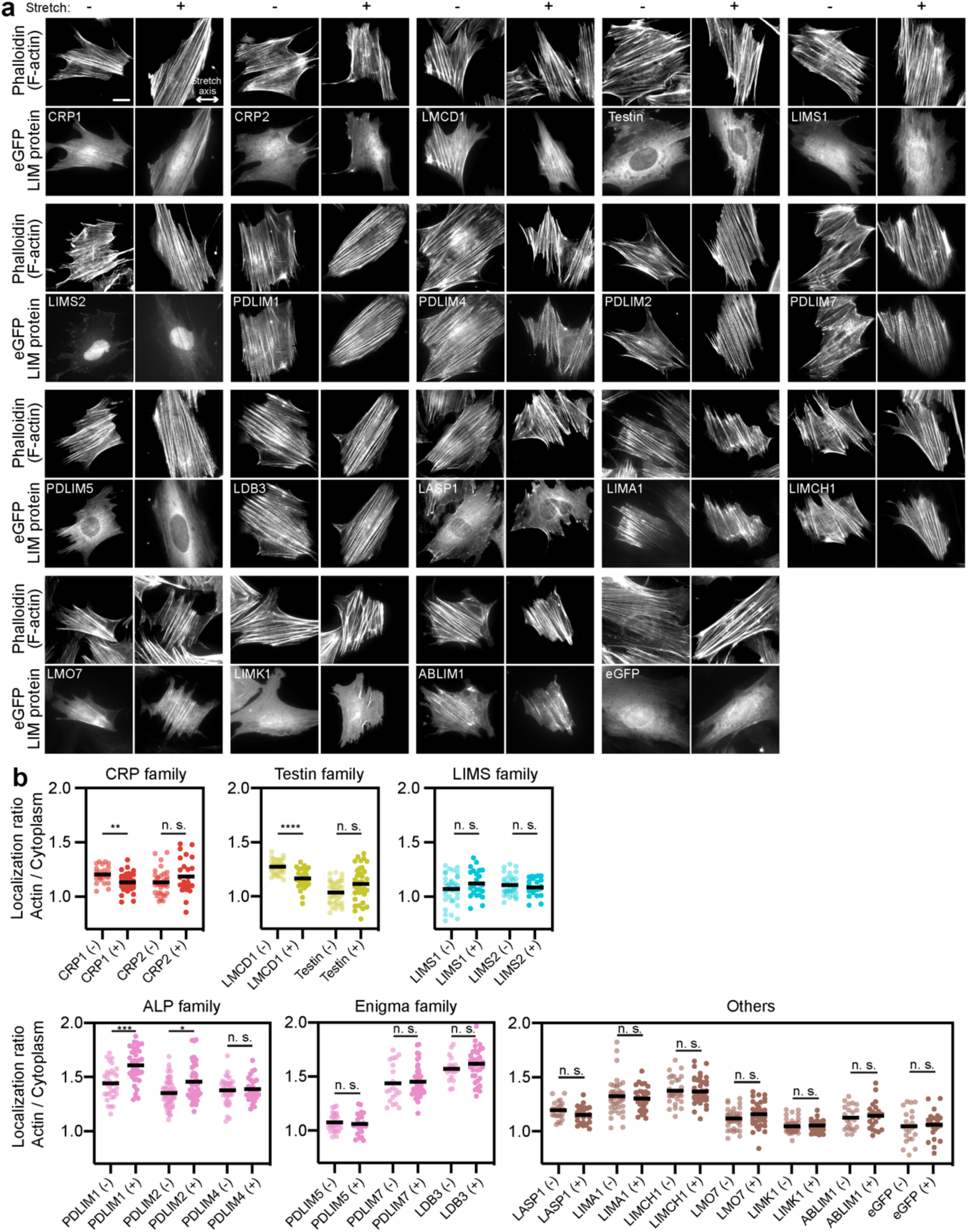
**Non-mechanoaccumulating LIM proteins identified in the cell-stretching screen. a**, Epifluorescence micrographs of eGFP-labeled LIM proteins in unstretched (-) and stretched (+) MEFs stained with phalloidin to label F-actin. Double-headed arrow indicates the uniaxial stretch direction. Scale bar, 20 μm. **b**, Whole-cell actin enrichment of LIM proteins in unstretched (-) and stretched (+) MEFs (20 ≤ n ≤ 60). Bars represent means. Dunnett’s T3 multiple comparisons test after Welch’s ANOVA is performed by comparing the actin enrichments of each LIM protein in unstretched cells with those in stretched ones simultaneously for all the LIM proteins included in the screen. NS, p > 0.05; ** p < 0.01; *** p < 0.001; **** p < 0.0001.

**Extended Data Fig. 4.**
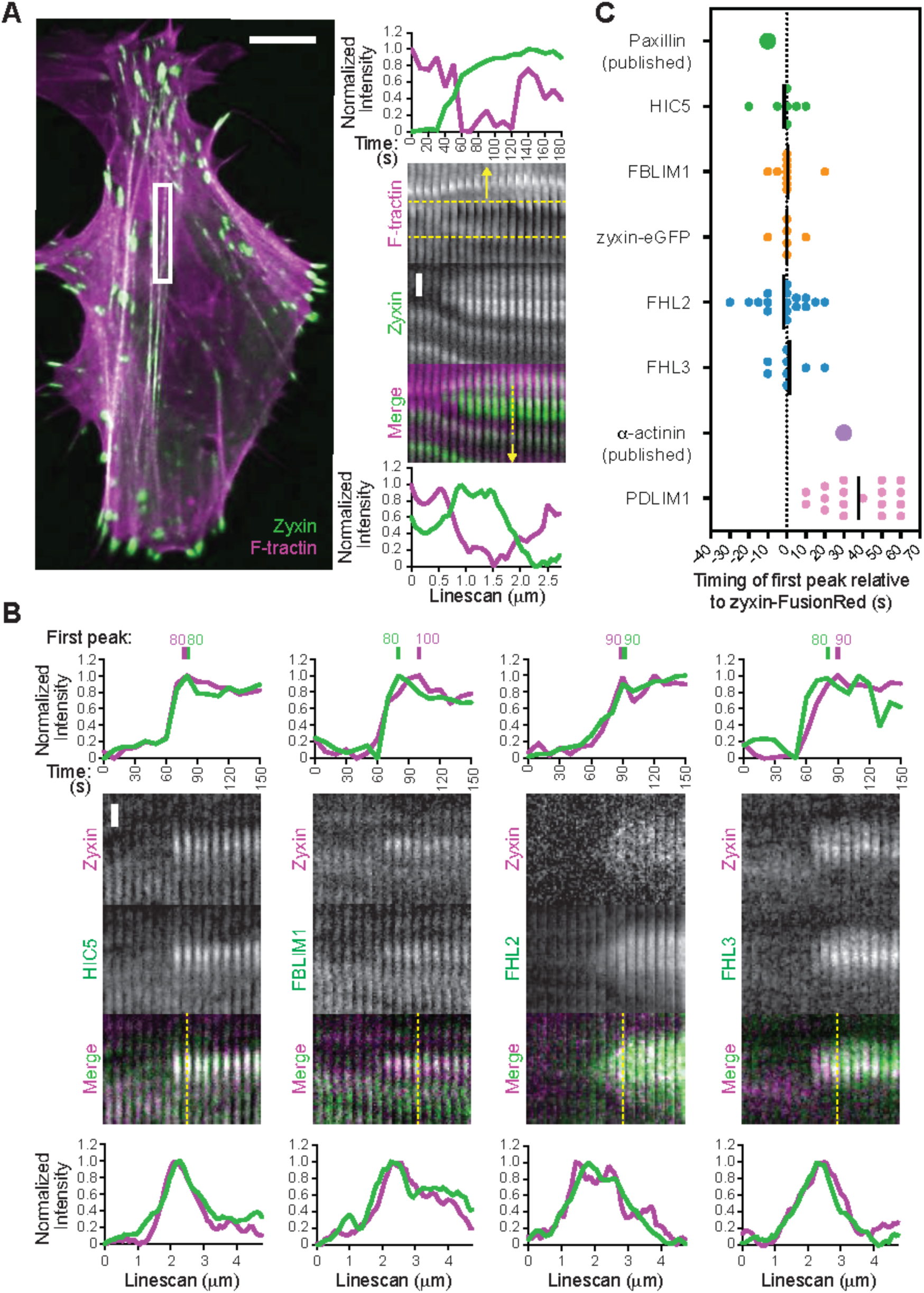
**FHL-, paxillin-, and zyxin-family proteins localize to SF strain sites with similar kinetics, supporting recognition of a common molecular mark. a**, Left, spinning disk confocal snapshot of a U2OS cell co-expressing F-tractin-mApple (magenta) and zyxin-eGFP (green). Scale bar, 5 μm. Box highlights an SF strain site. Right, intensity versus time plot (top), time-lapse montages (middle), and intensity line scan (bottom) of F-tractin and zyxin at an SF strain site. Scale bar, 1 μm. **b**, Intensity vs. time plots (top), time-lapse montages (middle), and intensity line scans (bottom) of representative SF strain sites marked by zyxin-FusionRed and the indicated eGFP-labeled LIM protein. Magenta, zyxin-FusionRed. Green, eGFP-labeled LIM protein. Scale bar, 1 μm. The time of first intensity peak is indicated for each protein. **c**, Timing of first intensity peak of indicated LIM proteins at 6 ≤ n ≤ 17 SF strain sites relative to zyxin-FusionRed. Bars represent means. Dash line indicates simultaneous recruitment of zyxin-FusionRed and the eGFP-labeled LIM protein.

**Extended Data Fig. 5.**
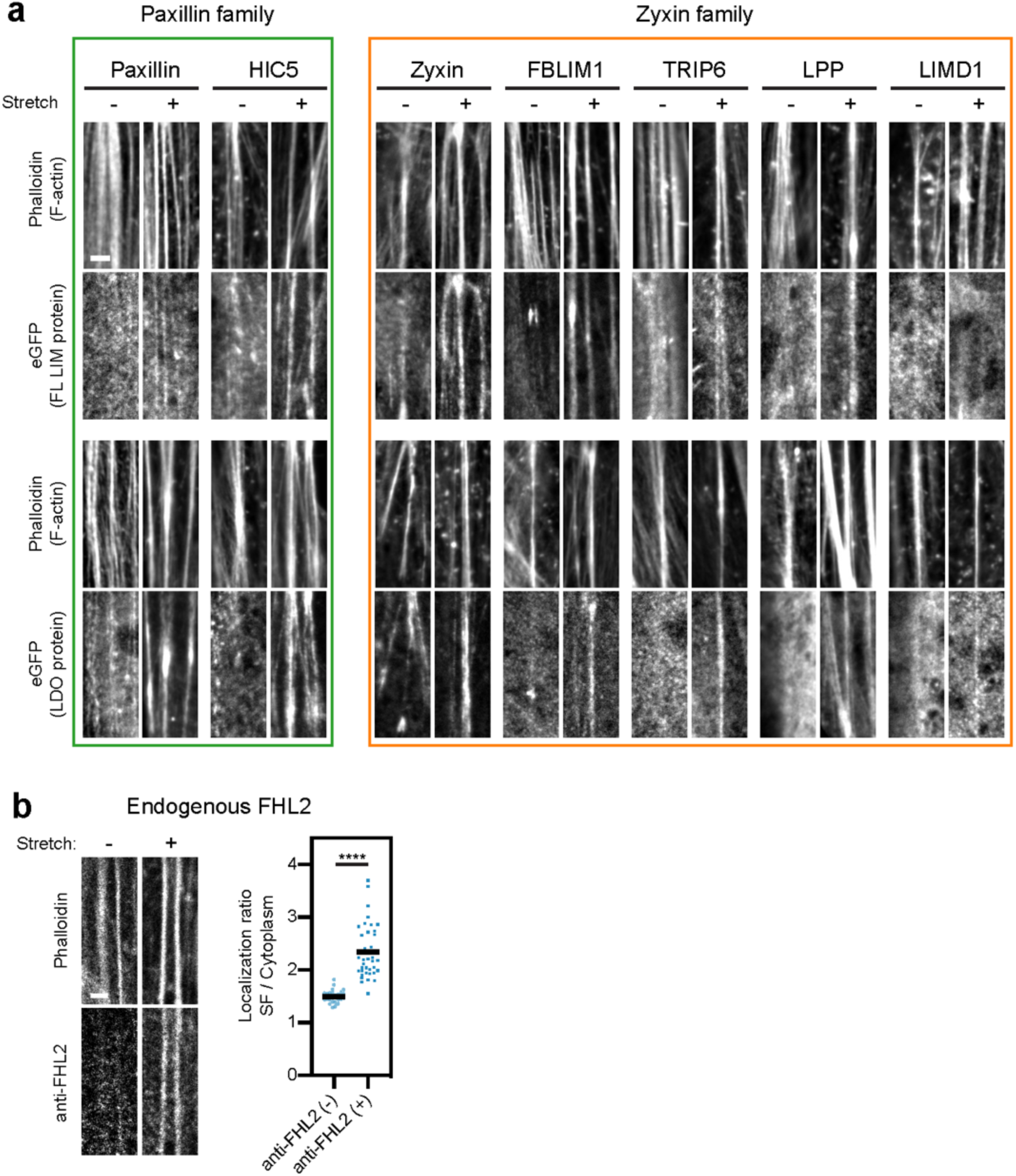
**Mechanoresponsive LIM proteins accumulate on SFs in stretched MEFs. a,** Confocal slices of phalloidin-stained SFs and eGFP-labeled paxillin- and zyxin-family proteins in unstretched (-) and stretched (+) MEFs. Top panel, full-length (FL). Bottom panel, LIM domain only (LDO). Scale bar, 2 μm. **b**, Left panel, confocal slices of SFs stained by phalloidin (top) and endogenous FHL2 (bottom) visualized by immunofluorescence. Scale bar, 2 μm. Right panel, SF enrichment of endogenous FHL2 in unstretched (-) and stretched (+) MEFs. Each data point is obtained from a single SF (25 ≤ n ≤ 38, N = 2 biological replicates). Bars represent means. Welch’s unpaired t-test: **** p < 0.0001

**Extended Data Fig. 6.**
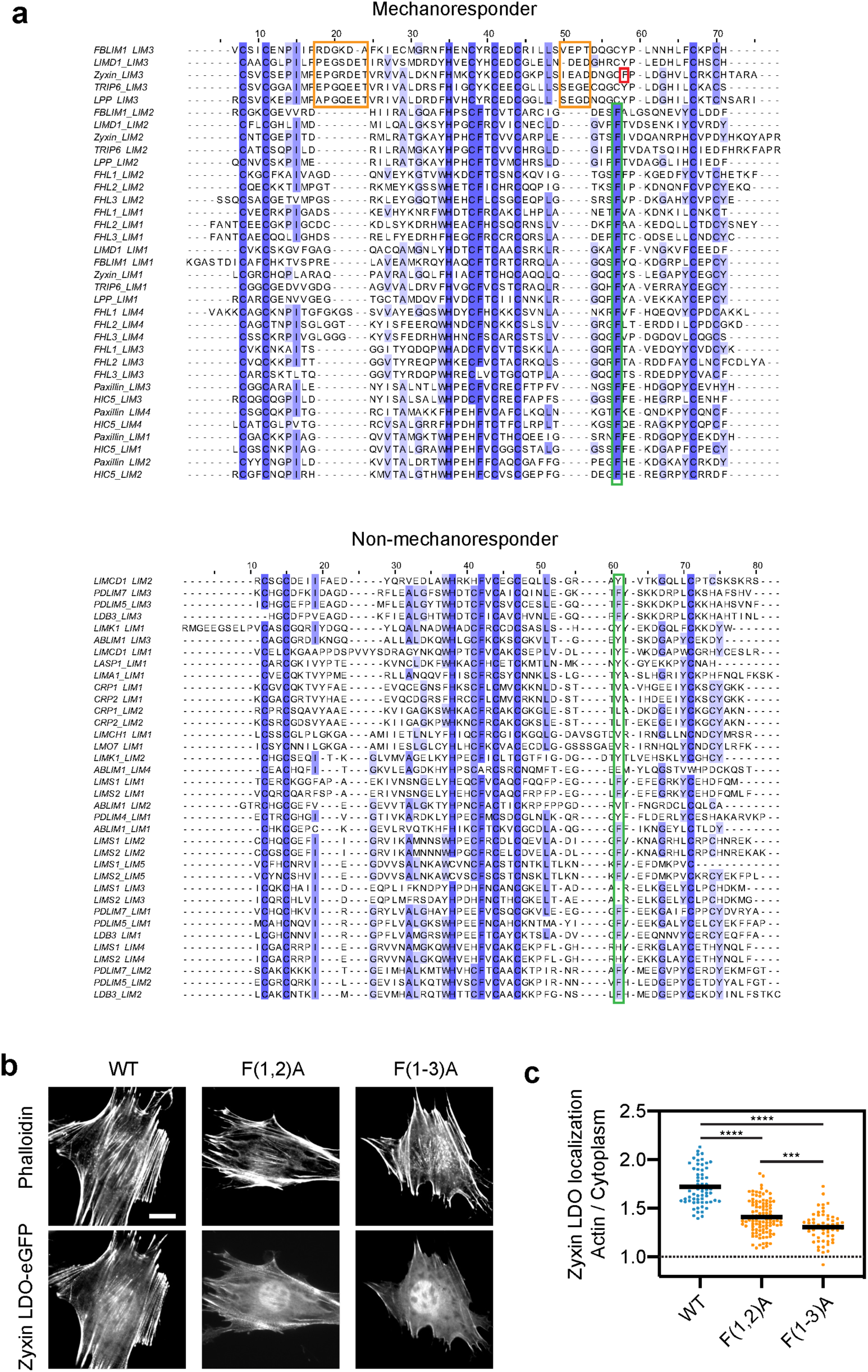
**A phenylalanine in the divergent LIM3 of zyxin additively contributes to mechanoaccumulation. a**, Multiple sequence alignments of mechanoresponder (top) and non-mechanoresponder (bottom) LIM domains, colored by conservation (shades of purple). Green boxes highlight the conserved phenylalanine in mechanoresponder LIM domains (top) and residues at the same position in non-mechanoresponder LIM domains (bottom). Red box highlights the phenylalanine next to the conserved position in zyxin LIM3. Orange boxes highlight extended insertions in LIM3 of zyxin-family members. **b**, Epifluorescence micrographs of MEFs expressing eGFP-labeled WT zyxin LDO (left), zyxin LDO F(1,2)A (middle), and zyxin LDO F(1-3)A (right), stained with phalloidin to label F-actin. Scale bar, 20 μm. **c**, Whole-cell actin enrichments of wild-type (n = 62), F(1,2)A (n = 102), and F(1-3)A (n = 56) zyxin LDO. N = 2 biological replicates. Games-Howell’s multiple comparison test after Welch’s ANOVA: *** p < 0.001; **** p < 0.0001.

**Extended Data Fig. 7.**
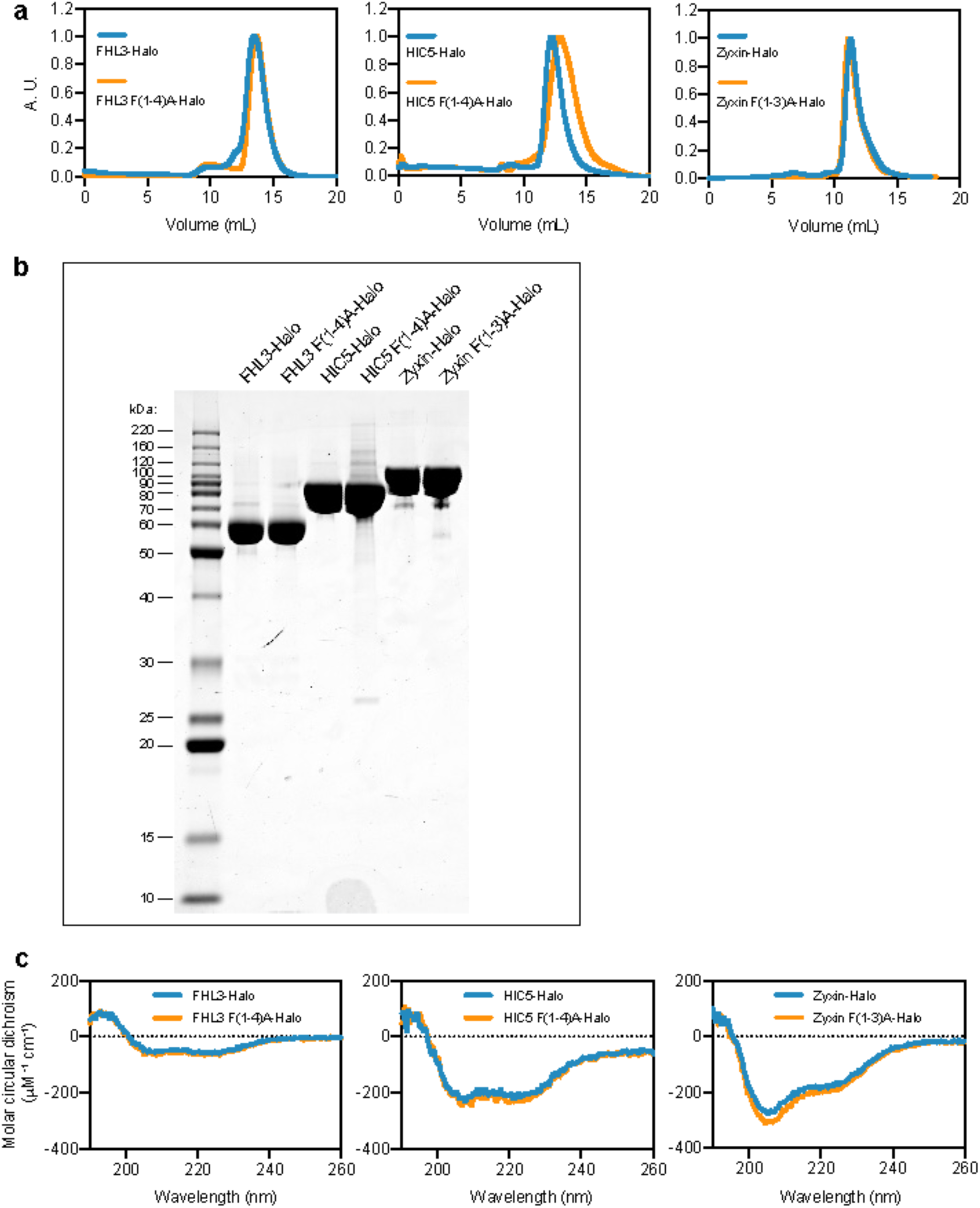
**Purified LIM proteins featuring phenylalanine mutations preserve their folded structures. a**, Normalized size-exclusion chromatograms of Halo-tagged wild-type and mutant FHL3 (left), HIC5 (middle), and zyxin (right). **b**, Coomassie-stained SDS-PAGE (3 μg) and **c**, circular dichroism spectra of the indicated purified proteins.

**Extended Data Fig. 8.**
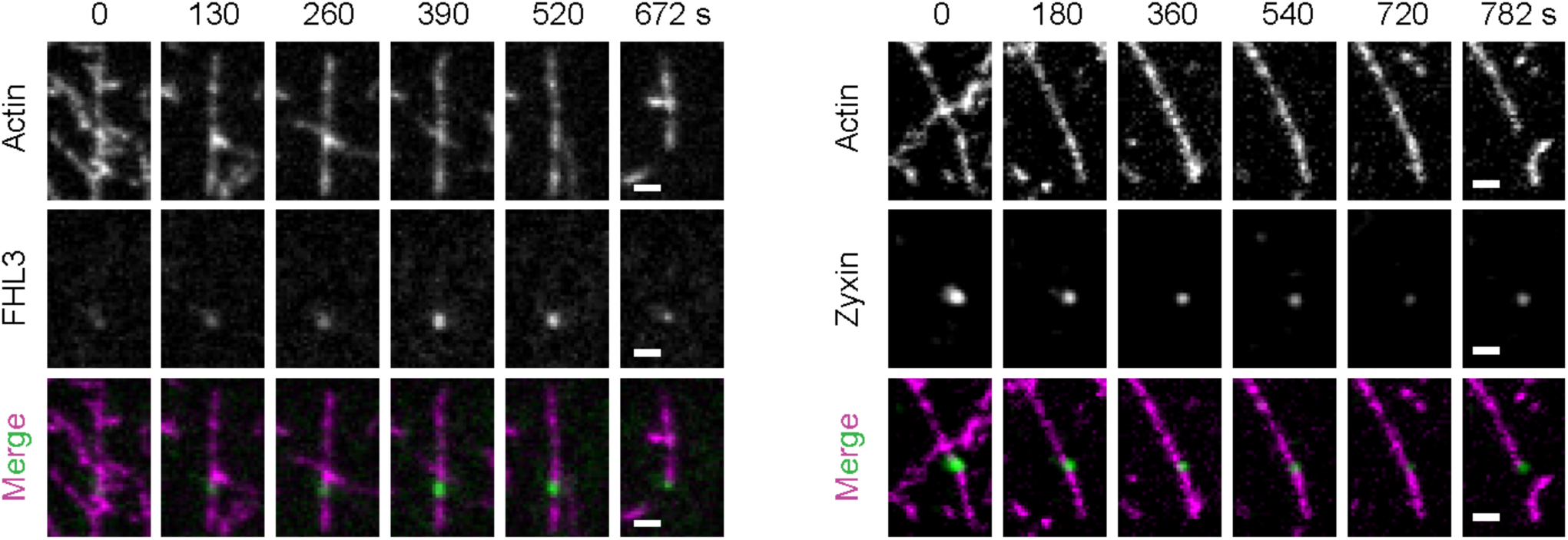
**LIM proteins rebinding after filament breakage.** Montages showing the rebinding of FHL3 (left) and zyxin (right) after filament breakage. Scale bars, 2 μm.

**Extended Data Fig. 9.**
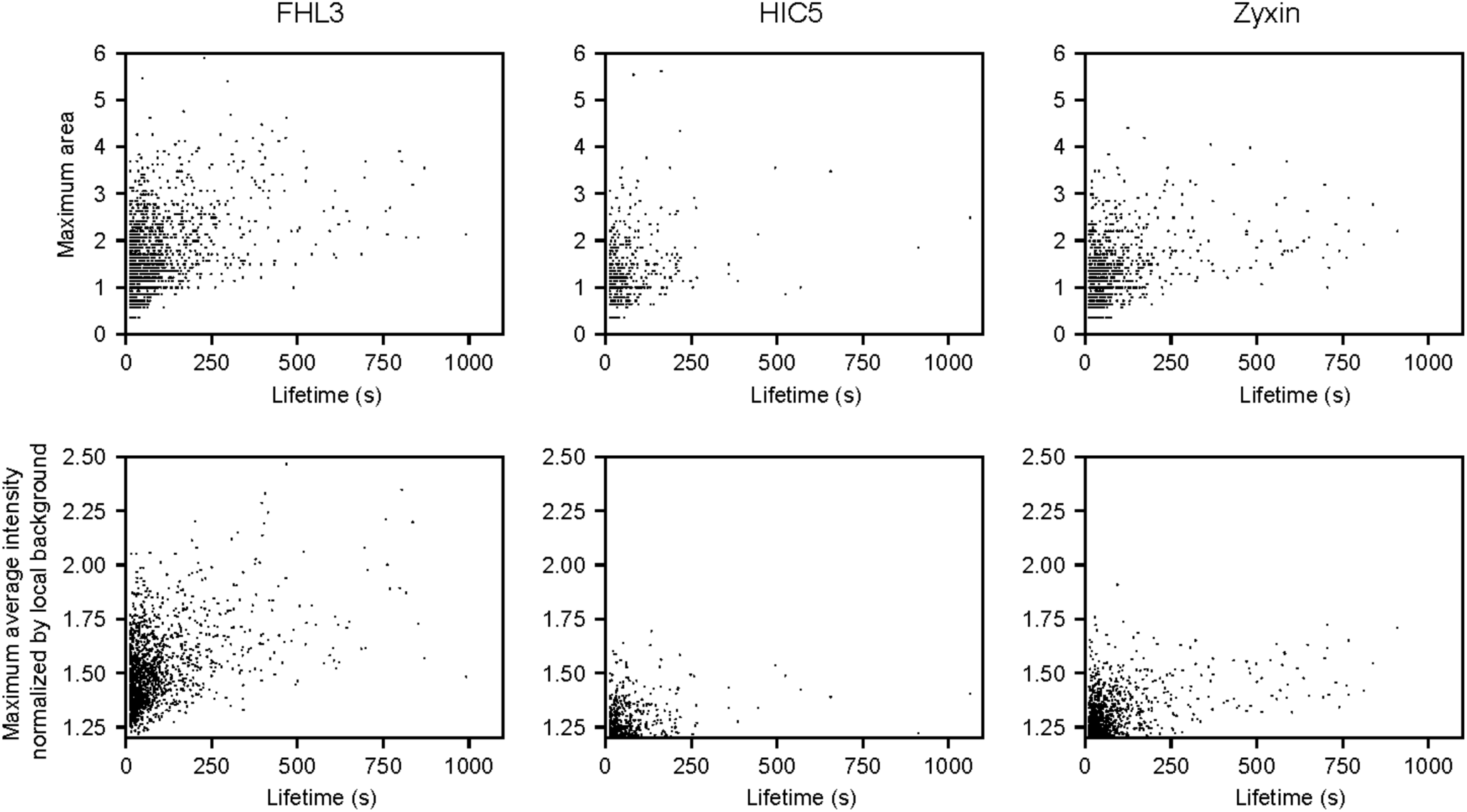
**Additional quantification of LIM protein patches.** Scatter plots of maximum area (top) and maximum normalized average fluorescence intensity (bottom) of patches versus their lifetime for the indicated proteins.

**Extended Data Fig. 10.**
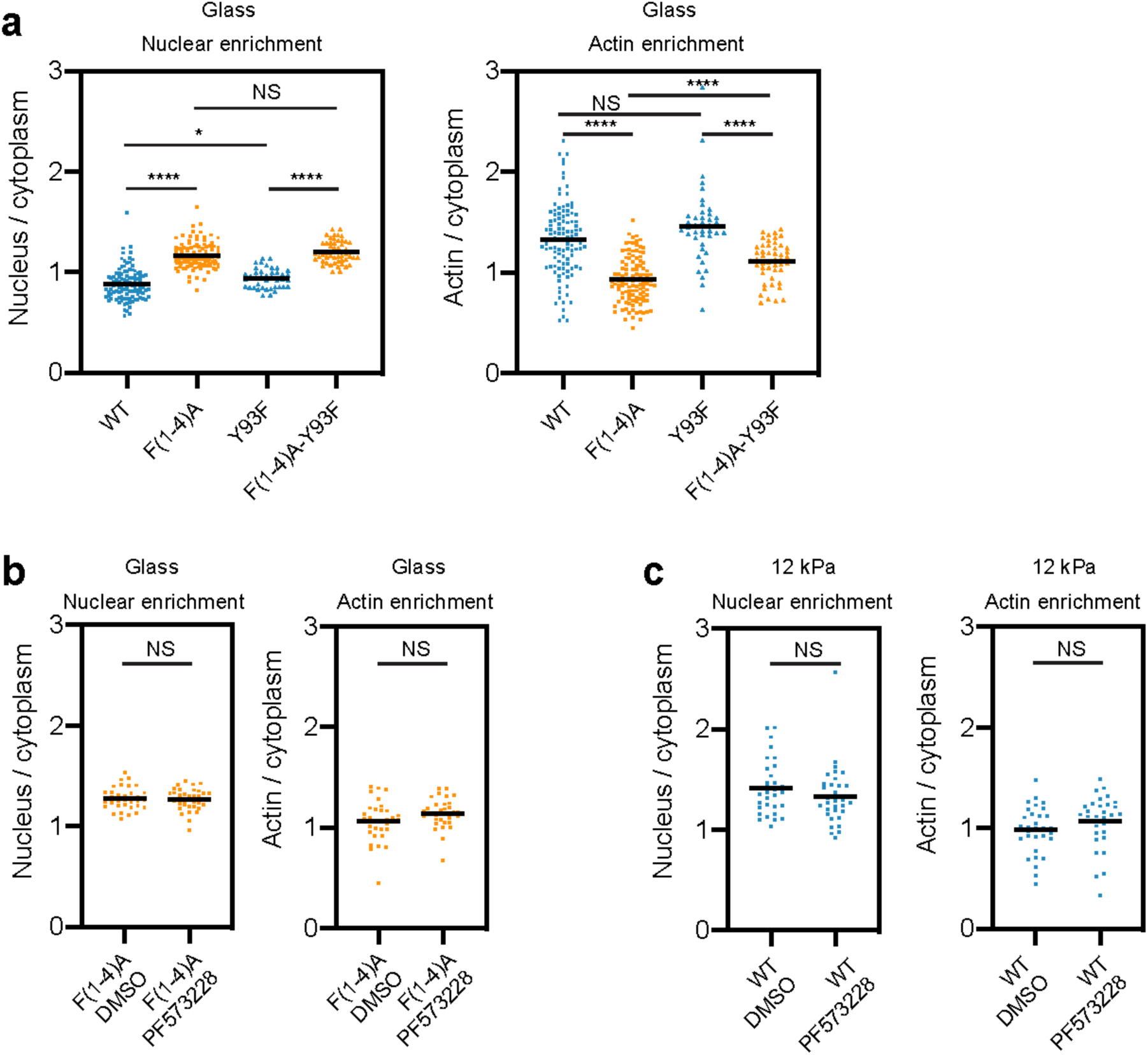
**FAK phosphorylation is not required for FHL2 nuclear shuttling in MEFs. a**, Nuclear (left) and actin (right) enrichment of eGFP-tagged wild-type (n = 106), F(1-4)A (n = 110), Y93F (n = 41), and F(1-4)A-Y93F FHL2 (n = 50) expressed in MEFs plated on glass. N = 2 biological replicates. Games-Howell’s multiple comparison test after Welch’s ANOVA: NS, p > 0.05; * p < 0.05; **** p < 0.0001. **b**, Nuclear (left) and actin (right) enrichment of eGFP-tagged FHL2 F(1-4)A expressed in MEFs plated on glass with (n = 34) and without (n = 36) FAK inhibition. N = 2 biological replicates. Welch’s unpaired t-test: NS, p > 0.05. **c**, Nuclear (left) and actin (right) enrichment of eGFP-tagged wild-type FHL2 expressed in MEFs plated on 12-kPa hydrogel with (n = 31) and without (n = 32) FAK inhibition. N = 2 biological replicates. Welch’s unpaired t-test: NS, p > 0.05.

**Extended Data Table 1.** Number of data points and outliers identified by ROUT method in GraphPad Prism for SF enrichment quantification in unstretched (-) and stretched (+) MEFs.

|  | Number of data points | Number of outliers |
| --- | --- | --- |
| **FHL1 (-)** | 28 | 0 |
| **FHL1 (+)** | 80 | 1 |
| **FHL2 (-)** | 69 | 5 |
| **FHL2 (+)** | 142 | 4 |
| **FHL3 (-)** | 166 | 9 |
| **FHL3 (+)** | 226 | 2 |
| **Paxillin (-)** | 72 | 2 |
| **Paxillin (+)** | 134 | 2 |
| **HIC5 (-)** | 102 | 1 |
| **HIC5 (+)** | 161 | 1 |
| **Zyxin (-)** | 74 | 5 |
| **Zyxin (+)** | 140 | 3 |
| **FBLIM1 (-)** | 91 | 2 |
| **FBLIM1 (+)** | 132 | 1 |
| **TRIP6 (-)** | 50 | 2 |
| **TRIP6 (+)** | 89 | 4 |
| **LPP (-)** | 39 | 2 |
| **LPP (+)** | 101 | 3 |
| **LIMD1 (-)** | 28 | 1 |
| **LIMD1 (+)** | 37 | 2 |
| **Paxillin LDO (-)** | 198 | 14 |
| **Paxillin LDO (+)** | 244 | 1 |
| **HIC5 LDO (-)** | 111 | 2 |
| **HIC5 LDO (+)** | 170 | 3 |
| **Zyxin LDO (-)** | 71 | 3 |
| **Zyxin LDO (+)** | 115 | 2 |
| **FBLIM1 LDO (-)** | 100 | 2 |
| **FBLIM1 LDO (+)** | 146 | 0 |
| **TRIP6 LDO (-)** | 45 | 1 |
| **TRIP6 LDO (+)** | 69 | 1 |
| **LPP LDO (-)** | 129 | 3 |
| **LPP LDO (+)** | 222 | 2 |
| **LIMD1 LDO (-)** | 35 | 2 |
| **LIMD1 LDO (+)** | 93 | 2 |

## Video captions

**Supplementary Video 1 |** Spinning-disk confocal live-cell imaging of SF strain sites in MEFs co-expressing F-tractin-mApple and the indicated eGFP-labeled constructs. Scale bar, 3 μm. Time is minutes : seconds.

**Supplementary Video 2 |** TIRF imaging of *in vitro* force reconstitution assay in the absence (left) and presence (right) of ATP. Scale bar, 10 μm. Time is minutes : seconds.

**Supplementary Video 3 |** FHL3 dissociates from an actin filament upon filament failure, and rebinds to the filament as load develops. Scale bar, 2 μm. Time is minutes : seconds.

**Supplementary Video 4 |** FHL3 patches bind to the middle and to the end of actin filaments, and unbind upon filament breakage. Scale bar, 2 μm. Time is minutes : seconds.

**Supplementary Video 5 |** An FHL3 patch elongates along a tensed actin filament, and unbinds upon filament breakage. Scale bar, 2 μm. Time is minutes : seconds.

**Supplementary Video 6 |** Spinning-disk confocal live-cell imaging of a MEF co-expressing LifeAct-mApple and FHL2-eGFP pre-treated with Hoechst to label the nucleus. 250 nM cytochalasin D was added at 10 minutes and washed out 60 minutes after the time-lapse imaging started. Scale bar, 20 μm. Time is minutes : seconds.

